# Investigating Spatiotemporal Dynamics of Cortical Activity During Language Production in the Healthy and Lesioned Brain

**DOI:** 10.1101/2023.04.27.538530

**Authors:** Quentin Mesnildrey, Alexandre Aksenov, Malo Renaud D’Ambra, Gesa Hartwigsen, Vitaly Volpert, Anne Beuter

## Abstract

Efficient language production requires rapid interactions between different brain areas. These interactions can be severely affected by brain lesions. However, the neurophysiological correlates of the spatiotemporal dynamics during language production are not well understood. The current pilot study explores differences in spatiotemporal cortical dynamics between five subjects with post-stroke aphasia and five control subjects. Electroencephalography was recorded during picture naming in both groups.

Average-based analyses (event-related potential (ERP), frequency-specific Global Field Power (GFP)), reveal a strong synchronization of cortical oscillations, especially within the first 600ms post-stimulus, with a time shift between participants with aphasia and control subjects. ERPs and the corresponding brain microstates indicate coordinated brain activity alternating mainly between frontal and occipital zones. This behavior can be described as standing waves between two main sources.

At the single-trial scale, traveling waves (TW) were identified from both phase and amplitude analyses. The spatiotemporal distribution of amplitude TW reveals subject-specific organization of several interconnected hubs. In patients with aphasia this spatial organization of TW reveals zones with no TW notably in the vicinity of stroke lesions.

The present results provide important hints for the hypothesis that TW contribute to the synchronization and communication between different brain areas especially by interconnecting cortical hubs. Moreover, our findings show that cortical dynamics is affected by brain lesions.

**Contribution to the Field:**

1. Spatiotemporal cortical dynamics of individual trials reveals the presence of phase and amplitude traveling waves.
2. Exploration of traveling waves on the 2D cortical surface reveals the presence of interconnected epicenters or hubs in all subjects.
3. The spatiotemporal distribution of traveling waves shows a higher density in the prefrontal area for people with aphasia than for healthy subjects.
4. For subjects with aphasia, a sparser density of traveling waves is observed in the approximated lesion area.
5. Event-related potential analyses reveal a consistent alternating activity between the frontal and occipital regions.
6. Subjects with aphasia present a larger and/or delayed contribution in the delta range in the GFP patterns compared to control subjects.

## 1. Introduction

Language is organized in distributed networks in the human brain. Across the last decades, our knowledge about the functional organization and specialization of language networks has considerably improved. Neuroimaging and lesion studies have identified key areas for language comprehension and speech production and point towards both overlap and specialization of networks for specific linguistic functions (Price, 2000, 2010; Vigneau et al., 2011; Hodgson et al., 2021). Moreover, recent research provides insight into the spatial dynamics of language reorganization after brain lesions (Saur et al., 2006; Stockert et al., 2020). However, less is known about the spatiotemporal dynamics of language network organization in the healthy and lesioned brain. A better understanding of such processes may help to refine current models of language (re-)organization and ultimately improve current treatment approaches for people with speech and language impairment after brain lesions.

To further improve our understanding of language network organization, one needs to clarify the link between spatial network organization and temporal dynamics during language processing. Relating these dynamics to specific language operations will help better understand its relevance at the behavioral level. In parallel, identifying differences in the spatiotemporal dynamics between healthy and lesioned brains is of major importance.

Decades of research in the field of language production have enabled the description of the spatiotemporal dynamics of cortical processing in finer details (Vigneau et al., 2006; Llorens et al., 2011; Pham et al., 2013; Hartwigsen, 2015; Liljeström et al., 2015; Klaus et al., 2019; Piai et al., 2019; Sarubbo et al., 2020; Ala-Salomäki et al., 2021) and several models of cortical processing have been proposed (Strijkers et al., 2010; Indefrey, 2011; Laganaro, 2017). Classical models of speech production generally contain two lexical processing stages, a lexical semantic stage in which the meaning of a word is accessed, and a lexical phonological stage in which its sound code is accessed (e.g., Rapp & Goldrick, 2006; Levelt, Roelofs, & Meyer, 1999; Griffin & Bock, 1998; Caramazza, 1997; Dell et al. 1997; see Graves et al., 2007). Existing models differ with respect to the degree of discreteness versus seriality of information flow during speech production (Graves et al., 2007). Serial models claim that lexical semantic access must be completed before lexical phonological access can begin to take place (e.g., Levelt et al., 1999). In contrast, cascade models propose that lexical phonological access can begin before lexical semantic access is complete (e.g., Humphreys, Riddoch, & Quinlan, 1988). Finally, interactive models propose that processing at the lexical phonological level can feed back to the lexical semantic level (e.g., Dell et al., 1997). Despite these differences in information flow dynamics, the different models agree that during picture naming, lexical semantic information flow normally begins to be accessed prior to lexical phonology (see Graves et al., 2007). Accordingly, one may summarize the main steps of the picture naming process as follows: visual processing, visual recognition, semantic processing, lemma retrieval, phonological encoding, articulation programming, and speech production, in parallel with self-monitoring (e.g., Indefrey and Levelt, 2004; Dell, Martin and Schwartz, 2007).

With respect to the underlying neural substrates of language, several key areas have been identified for lexical encoding (e.g. posterior middle temporal gyrus), semantic processing (e.g. angular gyrus, anterior inferior frontal gyrus), or phonological processing (e.g. inferior frontal gyrus, supramarginal gyrus), (Devlin et al., 1998; Vigneau et al., 2006; Moliadze et al., 2019; Stockert et al., 2020). However, despite the spatial location of these areas, the mechanisms of their synchronization and/or sequential activation, their functional and/or anatomical relationship with homologue regions and neighbor brain areas remain unclear and variability among studies is important (Hampshire et al., 2010; Mehrkanoon et al., 2014; Hartwigsen et al., 2020, 2021). Consequently, the dynamics of such network interactions remain largely unexplored. In particular, it remains unclear how brain lesions affect network interactions and whether a lesion changes the spatiotemporal dynamics of language production.

Across the last decades, several studies have shown that cortical oscillations propagate at the surface of the human cortex forming so-called traveling waves (TW, Massimini et al., 2004; Muller et al., 2018b; Zhang et al., 2018; Alamia et al., 2019). Such waves are described as a spatiotemporally coherent evolution of either the phase or the amplitude of electrophysiological signals (sometimes referred to as phase waves and amplitude waves). Complementary studies attempted to clarify the functional role of these TW (Alexander et al., 2013; Muller et al., 2018; Ermentrout & Kleinfeld, 2001; Beuter et al., 2020; Davis et al., 2020). It seems, that TW per se do not carry the information from one region to another but rather contribute to the synchronization, optimization of network connectivity and to the processing of sensory inputs. One may thus hypothesize that TW indirectly participate in the efficient treatment of neural signals. In other words, they appear to play an auxiliary but important functional role.

Considering such complex and high-level mechanisms, cortical damage resulting from various causes (e.g., stroke, trauma, and tumor) inevitably induces physiological disorders. Brain lesions induce local changes in the electrical conductivity of neural tissue. As shown by Bessonov et al., 2019, 2020, these changes may disturb or prevent the passage of TWs and thus yield a poorer connectivity which may eventually harm cortical processing and network interactions. These disorders affect executive or cognitive capacities with different degrees of severity depending on numerous factors. Such factors include the location of the lesion, its spatial spread and depth. Stroke lesions in particular disturb the entire connectome (Klingbeil et al., 2019). In response to these disturbances, complex plastic changes and reorganization are being observed either by an upregulation of domain general areas in the vicinity of the lesioned area or by the recruitment of contralateral homologous area depending on the location of the lesion (Hartwigsen, 2016; Hartwigsen et al., 2020, 2021).

In the present pilot study, we recorded the electroencephalogram (EEG) in healthy subjects and people with post-stroke aphasia performing a picture naming task. The first aim of this work is to investigate the spatiotemporal cortical dynamics by means of both static and dynamic methods. A secondary aim of this study is to characterize regular and pathological behaviors to assess the influence of stroke lesions as well as plasticity induced changes on cortical activity.

## 2. Materials and Methods

### 2.1. Participants

Ten adult subjects took part in this pilot study. Five of them were healthy subjects and the other five subjects suffered from non-fluent post-stroke aphasia. The two groups were matched for age and gender (see details in table 1). All subjects were right-handed, assessed using the Edinburgh Handedness Inventory (Oldfield, 1971, see scores in table 1). Participants with aphasia (labelled AXX) were all diagnosed with Broca’s aphasia and a moderate degree of dysarthria resulting from a first-ever ischemic stroke of the left middle cerebral artery affecting the sylvian area. They were all in chronic phase, at least six months post stroke. While these criteria were rather restrictive, the exact location of the lesioned brain area as well as the severity of their language and/or motor disorders varied across subjects. A02, A04, and A05 suffered from severe motor deficits while A01 and A03 did not. The entire protocol was approved by the local ethics committee (Institutional review board, IRB-Euromov, number 2111D), all tests were conducted with the participants’ written informed consent and participants were paid for their involvement. All subjects were screened for normal cognitive functions using the Mini-Mental State Examination (score> 20, Derouesné et al., 1999; Kalafat et al., 2003). The Language Screening Test (Flamand-Roze et al., 2011) was realized by participants with aphasia in order to provide a quick and objective measure of the type and degree of their aphasia prior to testing without diagnostic purposes.

**Table 1.** Subjects’ details. From left to right: Subject number, age at testing, gender, duration since stroke, naming task score (%), Edinburgh test, screening test, cognitive test outcome measures and lesion information.

| Subject | Age<br>(years) | sex | Duration<br>since<br>stroke<br>(years) | Naming<br>score<br>(n=100) | Right<br>Handedness<br>dominance<br>(%) | Aphasia<br>screening<br>test | MMSE | Stroke lesion |
| --- | --- | --- | --- | --- | --- | --- | --- | --- |
| C01 | 49 | M | n/a | 93 | 100 | n/a | 24 | n/a |
| C02 | 56 | M | n/a | 99 | 100 | n/a | 29 | n/a |
| C03 | 71 | F | n/a | 92 | 100 | n/a | 25 | n/a |
| C04 | 71 | M | n/a | 95 | 100 | n/a | 28 | n/a |
| C05 | 57 | F | n/a | 97 | 100 | n/a | 29 | n/a |
| A01 | 50 | M | 3 | 97 | 100 | 13 | 26 | Left frontoparietal<br>Deep and<br>superficial |
| A02 | 56 | M | 4.5 | 97 | 100 | 15 | 28 | Left frontoparietal,<br>Deep and<br>superficial |
| A03 | 70 | F | 4 | 88 | 89 | 13 | 22 | Left<br>temporoparietal,<br>posterior sylvian<br>area<br>Superficial |
| A04 | 71 | M | 20 | 88 | 100 | 13 | 27 | Left temporal,<br>Superficial |
| A05 | 56 | F | 3 | 89 | 100 | 15 | 26 | Left<br>frontotemporal,<br>Superficial |

All data were acquired during a single session lasting two hours. During these two hours the participant realized the aforementioned tests, then the experimenter installed the EEG system on the participant (see details in Section 2.3) and the participant realized the naming experiment. All tests were conducted in a quiet room.

### 2.2. Picture naming task

The participants sat approximately 60 centimeters away from a computer screen. A single experimenter sat outside their visual field and provided instructions to the participant.

The main task consisted in a free picture naming task. Each trial started with a blank screen for 1.5 seconds and subjects were allowed to breathe and blink normally. Subjects were then asked to fixate a white cross displayed at the center of the screen while limiting facial movements. After 1.5+/-0.2 seconds a black and white picture was displayed, and subjects had to name the recognized object. Subjects were asked to name the objects at their own pace and to provide one single response. For each trial, subjects’ oral response was recorded using an external microphone located halfway between the participant and the computer screen. The voice onset was automatically detected, and a marker was added accordingly to estimate the naming latency. The next trial was automatically launched 4s after the voice onset. The coherence of automatic voice detection (thresholding) and the actual audio signal was manually verified offline and corrected when necessary (e.g., background noise, hesitation, or multiple responses).

Pictures were selected from the Snodgrass corpus (Alario et al., 1999). The set of 100 images was divided in two runs of 50 items with a small rest between them. Among the 100 images, three items were repeated 10 times each and were referred to as control items (“DOG”, “APPLE” and “BED”) the other 70 images were presented once. Only frequent nouns were included (familiarity > 1.3). To avoid ceiling effects, 8 out of 100 words with higher complexity were included (familiarity < 2, visual complexity > 4, see Alario & Ferrand, 1999). All items were presented in a randomized order.

During the entire procedure and for each trial, several markers were added to the data. Markers 1 to 4, respectively correspond to the beginning of the trial, the cross on the screen, the image presentation, and the voice onset. In case of incorrect, unclear response or failed voice detection, the corresponding trials were manually tagged by the experimenter. The tagged items were then retested at the end of the set. This procedure guarantees a correct acquisition of all items.

Four minutes of resting state recording was performed before and after the picture naming task. For the first two minutes, participants were instructed to look at a white fixation cross in the middle of the computer screen (eyes-open condition), and then close their eyes for the last two minutes (eyes-closed condition).

### 2.3. Electroencephalogram

#### 2.3.1. Device and Software

All recordings were made using the Starstim-32 system (Neuroelectrics®). Attention was given to prevent acoustic and electromagnetic noise during the EEG acquisition. Continuous EEG recordings were obtained via 30 contacts (NG Geltrode) placed on the scalp using the electrode configuration depicted in Figure 1.A at a sampling frequency of 500 Hz. Two additional electroocculography electrodes (EOG) were positioned above and below the right eye to detect eye-related artifacts. The reference electrodes (CMS and DRL) were clipped on the right ear lobe. The protocol was designed using NIC2 software (Neuroelectrics, Barcelona, Spain), PsychoToolbox (Brainard, 1997) and custom Matlab interfaces (Matlab 2021, the Mathworks, Natick, CA).

**Figure 1.**
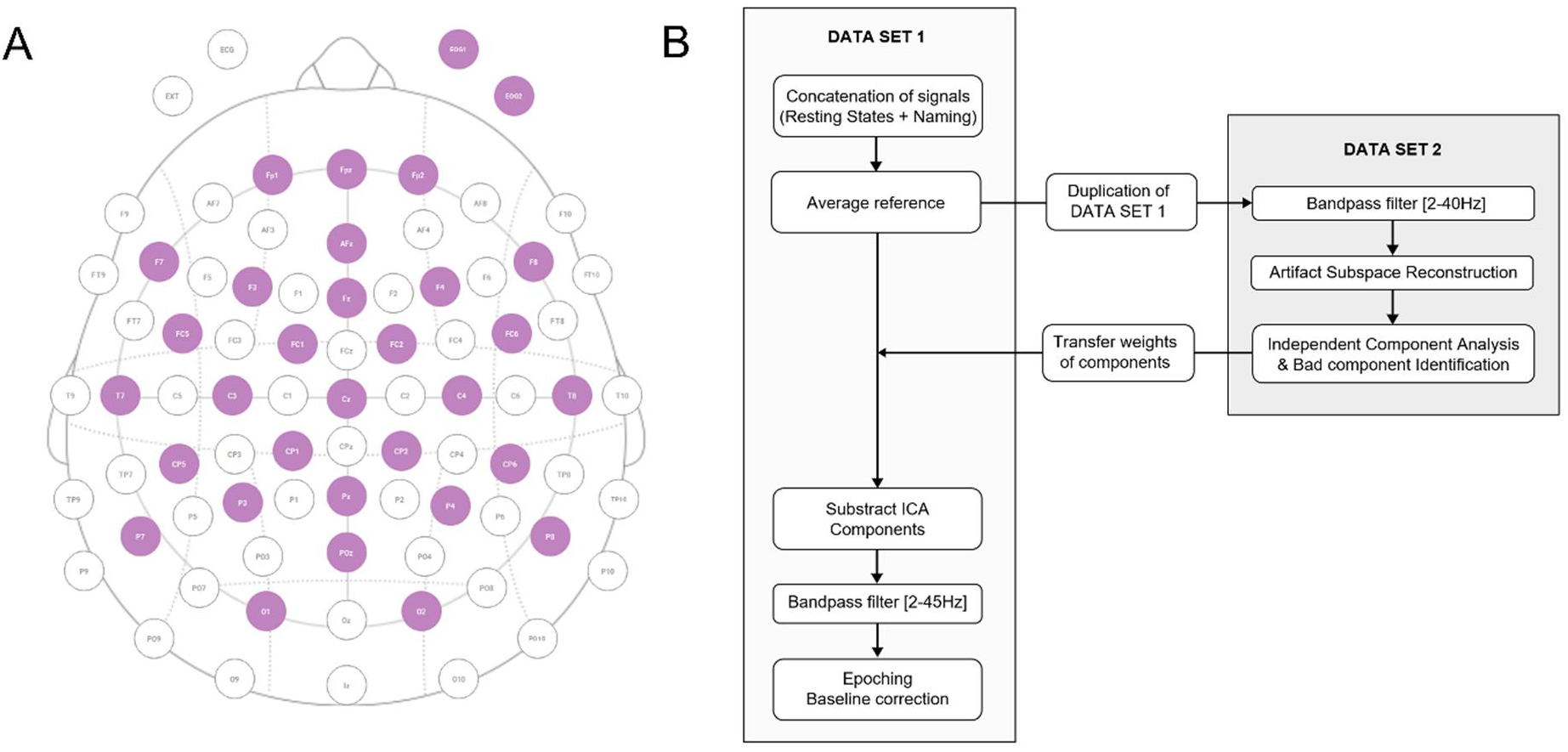
**(A)** Electrode configuration used for EEG recordings. Purple circles represent 30 active scalp contacts + 2 EOG contacts. **(B)** Preprocessing pipeline.

#### 2.3.2. Preprocessing

Off-line signal processing and analyses were performed using EEGLAB open-source toolbox (Delorme et al., 2004) and custom Matlab scripts. Raw and continuous EEG signals were preprocessed using the pipeline described in figure 1.B (EEGLAB functions). The relevance of independent component analysis (ICA) components was assessed, and bad components were removed. Movement and muscle artifacts were also visually identified and removed when necessary. The lowest cut-off frequency of the bandpass filters was fixed at 2Hz to remove slow drifts occurring in several datasets (C01, C02, and A05).

The different trials (n=100+retest) were isolated from the continuous data from 1.5s before image presentation to 4s after image presentation, thus resulting in 5.5s-long epochs. Note that while most items could be named within this default response window, few trials required more time. These trials were analyzed separately, and the actual number of regular epochs is mentioned in further analysis.

### 2.4 Analysis

EEG signals were analyzed using different approaches referred to as *static* and *dynamic*. The static methods included the calculation of frequency spectra as well as event-related techniques.

The power spectral density (PSD) was estimated from the epoched data (n=100) using Welch’s overlapped segment averaging estimator, with Hamming windows of length 5s and overlap of 2.5s (spec_topo of EEGLAB and custom Matlab codes). To estimate the peak frequency for each subject, the logarithmic aperiodic trend was first subtracted (Zhang et al., 2018; Donoghue et al., 2020). The peak frequency was then defined as the median of the individual

For each EEG channel, event-related potentials (ERPs) were calculated as the across-trials average. As a result, the amplitude of ERP signals provides information on the repeatability across trials. ERPs are used to investigate main trends relative to a specific event (typically the appearance or change in stimulus (see Murray et al., 2008, for a review) with a view of assessing the correlation between an objective measure and psychophysical and behavioral observations. In the following, stimulus-locked and response-locked ERPs were analyzed.

The global field power (GFP, Lehmann et al., 1980, 1984) is defined as the standard deviation across channels and may thus relate to localized cortical activation or deactivation. In the present study, GFP was calculated either for broadband signals (2-40Hz) or in commonly used brain rhythm frequency ranges, i.e., delta (2-4Hz), theta (4-8Hz), alpha (8-12Hz) and beta-1 (12-17Hz), beta-2 (17-22Hz), beta-3 (22-30Hz) and gamma (30-45Hz). Very low frequency components (<2Hz) in EEG data usually have much larger amplitudes than higher-frequency components. Including them drastically smooths the GFP curves. Here, filtering the data above 2Hz has the direct consequence of providing finer details in the GFP. Finally, both GFP and ERPs highlight timings where the topographic distribution of activity shows a specific pattern. Topographic maps were also computed to assess relevant cortical patterns (e.g., Chantsoulis et al., 2017; Laganaro, 2017; Mheich et al., 2021).

In the so-called dynamic approach several tools were designed to investigate the spatiotemporal cortical dynamics. Preprocessed EEG data were filtered with a 1Hz-wide band centered at nine different frequencies logarithmically spaced between 3 and 40 Hz (3, 5, 7, 9, 12, 16, 22, 30 and 40 Hz). The instantaneous phase of each filtered signal was obtained by computing the Hilbert transform. In order to assess the existence of phase waves along a specific line of electrodes we implemented a circular-linear regression (Aksenov and Beuter 2021; Fisher 1993; Zhang, et al. 2018).

The amplitude information was obtained using the following amplitude tracking technique. For each trial, the topographic activity map was computed for each sample (*topoplot* function) and used to locate the peak amplitude. This resulted in 2D trajectories of the peak amplitude. Additional treatments were applied on the trajectories to perform various analyses. In section 3.4, we first projected the 2D-trajectories extracted for each trial on a simple parceled scalp map (see Figure S4) and counted the frequency of occurrence of the maxima of amplitude in each of these different regions. This representation enables to evaluate the consistency of amplitude trajectories as well as the relative spatial distribution across trials and for each time step. Besides, contrary to ERP signals, it is less affected by the across-trial differences in amplitude since it only considers the location of the peak amplitude for each time frame and each trial.

The trajectories of amplitude were also represented in three dimensions to visualize their spatiotemporal evolution. We propose a method to perform an automatic segmentation of the trajectories in order to isolate specific patterns of cortical activity. From t=0 to the voice onset, we defined different polygons. Each polygon *Pi* was constructed as the projection on the scalp plane of the trajectory between *ti* and *ti+Tcycle*,where *Tcycle* is the period associated with the considered oscillating frequency. We calculated the Hausdorff distance (Hemanth, 2022) between successive cycles (*Pi-Pi+1*) with a time step of one cycle and the boundaries of each segment were then defined as the peaks of this distance. An empirical emergence threshold was applied to identify the main peaks of the Hausdorff distance.

In section 3.5 the following procedure was used to determine the existence of amplitude traveling waves. For each time step (sampling rate = 500Hz), the spatial distance between successive coordinates *i-1* and *i* was calculated, *Di-1, i*. If this distance did not exceed an empirical threshold, *Dmax*, samples *i-1* and *i* were attributed to the same wave. The end of a TW was determined as soon as *Di-1,i* exceeded *Dmax* or if *Di-1,i* was equal to zero (static activity). *Dmax* was defined considering a maximal wave velocity of 30m/s. Finally, we only considered TW lasting more than 8ms (i.e., four samples). Statistics and topographic maps of TW were then performed.

We also verified whether the TW identified using the present method could be detected using a phase-based approach. For each identified amplitude wave, we determined the closest EEG contact for each sample. We then limited our data to a subset of waves passing through (or next to) five distinct electrodes. Next, we extracted the instantaneous phase by computing the Hilbert transform of these five channels at a time corresponding to the middle of the TW. Finally, we fitted the phase values using a circular-linear regression method (Aksenov and Beuter 2021; Fisher 1993; Zhang, et al., 2018).

To investigate brain connectivity, we automatically located the main hubs revealed by TW topographic map. To do so, we stored the coordinates of both the source and destination of each TW detected using the previous algorithm. We then computed the density map of each set of coordinates using a square grid of 10-by-10 bins (Note that the default grid resolution of topographic representations is 67-by-67 pixels). For each dataset, main hubs were finally defined as the first ten maxima of the density map. Note that we did not consider hubs located at the edges of the interpolation grid since their accuracy is less dependable.

In the following, we display the findings from specific subjects. However, the same analyses were performed for all trials in all subjects. The corresponding figures are accessible in supplementary materials.

## 3. Results

### 3.1. Behavioral results

All participants were able to complete the protocol. The percentage of correct naming ranges from 92 to 99% for healthy subjects and between 88 and 97% for participants with aphasia. Depending on participants, false responses can be categorized as semantic paraphasia (“Fourchette” for “Cuillère” i.e., “Fork” for “Spoon”) potentially reflecting a lack of inhibition, phonemic paraphasia (phoneme substitution, addition, or anticipation. e.g., “thermoter” for “thermomètre” i.e., “thermoter” for “thermometer”), failure in memory retrieval, in particular for words that are known but not frequently used. In most failed trials it could be noticed that subjects with aphasia were aware of providing a false response. In a few other trials, they provided a correct answer while being convinced that they failed. Most participants with aphasia also showed different degrees of dysarthria. However, for these participants we consider these errors to result from a difficulty in associating phonetic and phonemic information rather than actual motor disorders (Blumstein et al., 1982; Kohn, 1988; Laganaro et al., 2010).

The naming latencies were also estimated from voice detection markers (median latency = 1530ms, sd = 3664ms for subjects with aphasia; median latency = 1376ms, sd = 1114ms for controls). The reasons for these relatively long naming latencies are discussed in Section 4. In few occurrences, participants with aphasia were not able to provide an answer at all or provided a wrong answer after a long period of time. The naming latency could thus not be estimated for these trials. In the following analysis, the exact number of considered epochs is stated. To evaluate whether subjects with aphasia required a longer period of time for the entire process of picture naming, we compare the naming latencies from successful trials only. A significant difference was found on the naming latency (for successful trials) between the two groups (t(933) = 3.50, p <0.001, mean difference = 215.5 ms). Figure S1 illustrates the distribution of naming latencies across groups.

### 3.2. Frequency spectrum

As expected, most spectra exhibited a clear maximum in the theta and alpha frequency bands except for C02 and C04. For subjects A01 and C02 the methods described by Zhang *et al*. failed to determine the peak frequency, the value was thus defined as the local maximum of the mean power spectral density for these two subjects. The corresponding values are reported in Figure 2. The mean peak frequency was 9.4 Hz (sd = 0.95Hz) and 8.6 Hz (sd = 2.24Hz) for healthy and participants with aphasia respectively. Note that A04 and A05 showed a peak frequency in the theta region. However, our small sample size did not yield any significant group effect.

**Figure 2.**
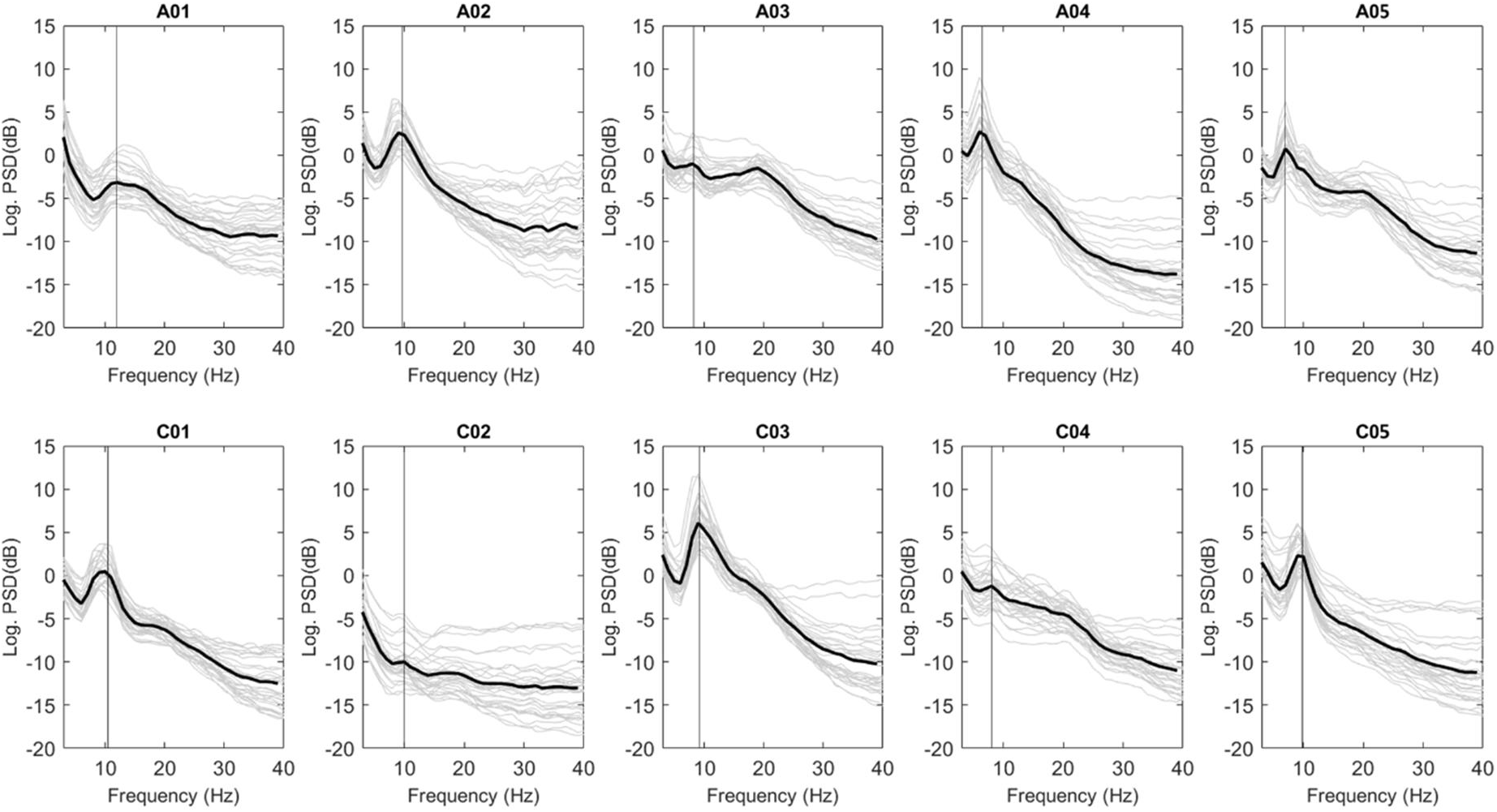
Power spectral density averaged across trials (n=100) for each participant (Cxx - healthy control participant, Axx - patient with aphasia). Grey lines represent power spectral density of individual electrodes, thick lines represent the average power spectral density. Vertical lines indicate the alpha peak frequency estimated after detrending.

### 3.3. Event Related Potential Analysis

#### 3.3.1. Global field power (GFP)

The average GFP calculated separately for both groups exhibited four main peaks located at 112, 176, 258 and 388ms post-stimulus for healthy subjects and at 120, 200, 264 and 410ms for participants with aphasia (see Figure 3.A). GFP patterns were consistent in both timing (four main peaks delayed by 20.5ms on average for subjects with aphasia) and amplitude for both groups across subjects.

**Figure 3.**
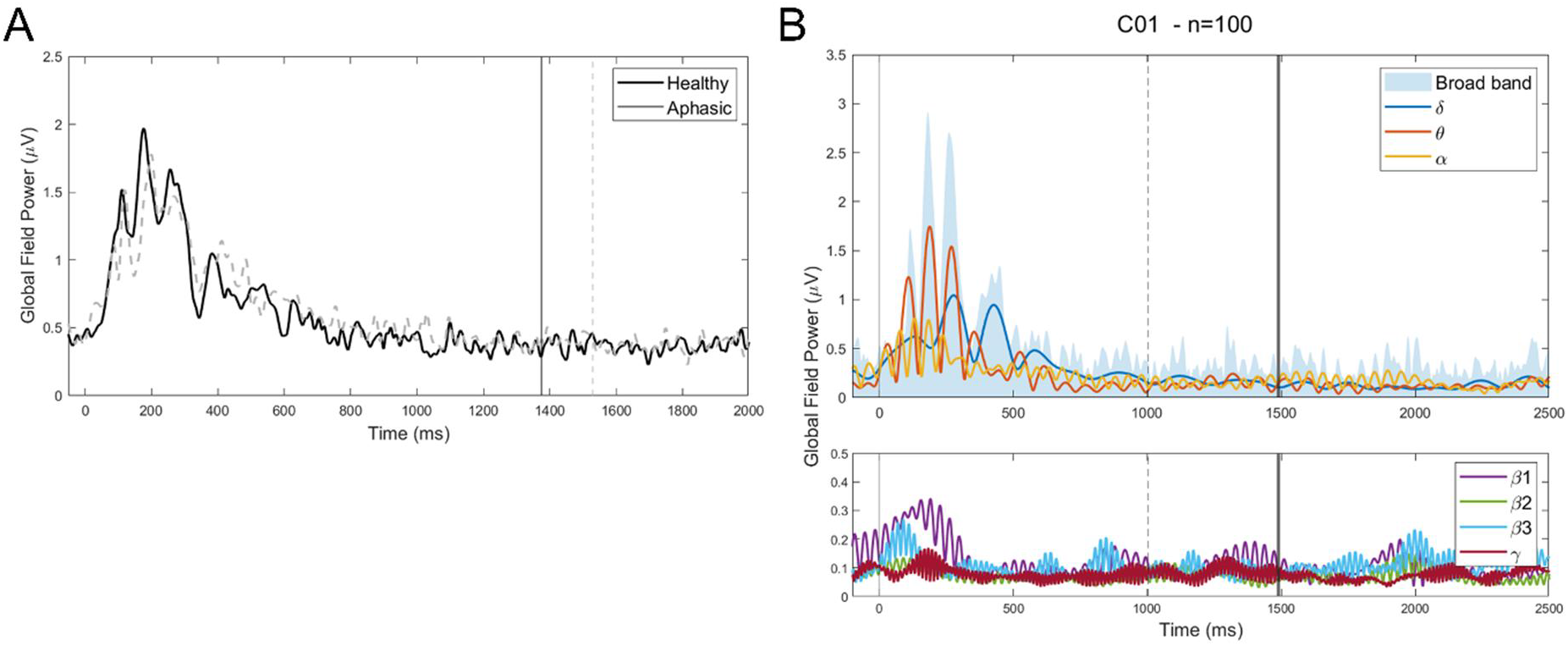
**(A)** Global field power for healthy subjects (black solid line) and participants with aphasia (dashed grey line). Data contain 100 epochs for each participant and were filtered between 2 and 40 Hz. Vertical lines indicate median naming latency. **(B)** Global field power in different frequency bands considering all trials from subject C01. High frequency ranges (Beta and higher) are displayed in the bottom subplot with an adapted scale for clarity. Vertical lines from left to right indicate the image presentation (t=0), the minimum, and median naming latency across the considered trials.

Next, we were interested in individual subject data. Individual GFP were thus calculated for all subjects. Figure 3.B shows a typical example of GFP calculated for subject C01 for broadband signals (2-40Hz) and in commonly used brain rhythm frequency ranges (see Section 2.4). Additional figures are provided in supplementary materials, Figure S2. Broadband data were very consistent with the group-level pattern (Figure 3.A), exhibiting between three and five main peaks of GFP within the first 600ms post-stimulus, depending on subjects.

In narrow frequency ranges, the largest values of GFP were also observed within the first 600ms. In the high frequency ranges (alpha and above) GFP patterns seem modulated by a slow component (∼1Hz). From a theoretical point of view, such a behavior can be obtained with the superposition of several oscillating sources with slight frequency differences (Volpert et al. 2022). It may also reflect the existence of frequency coupling (Lakatos et al., 2005; Klimesch, 2018). Potential implications for our understanding of the cortical dynamics are discussed below.

Of note, specific features of the broadband pattern of GFP resulted from the contributions of distinct frequency ranges. For all control subjects, the broadband GFP pattern was dominated by a theta component within the first 500ms. Moreover, in all control subjects, the first peak of GFP resulted from a theta and delta components, the second from theta and alpha components, while the third and fourth peak showed a large contribution of the delta component. This distribution of contributions was consistent across control subjects. In contrast, for subjects with aphasia, GFP patterns were much more heterogeneous. In particular, A01, A02, and A03 showed a larger contribution of the delta range relative to control subjects while A04 showed a delayed contribution of the delta component up to 1000ms (see Figure S2). This increased contribution of slow oscillations might reflect a pathological cortical activity as reported by several studies (Kamada et al., 1997; Meinzer et al., 2004; Spironelli et al., 2009).

To quantify the across-trials variability at the individual level we computed the GFP patterns for ten sets of twenty-five randomly selected epochs for each subject. We then measured the latencies of the four first peaks of GFP as well as the standard deviation across these ten iterations. At the individual level, a high across-trials consistency was found (see table 2). The most stable results were observed for the first peak post-stimulus in the theta and alpha bands. Overall, data from subjects with aphasia show a slightly poorer stability across the ten iterations (mean difference of 5.45ms across groups).

**Table 2.**
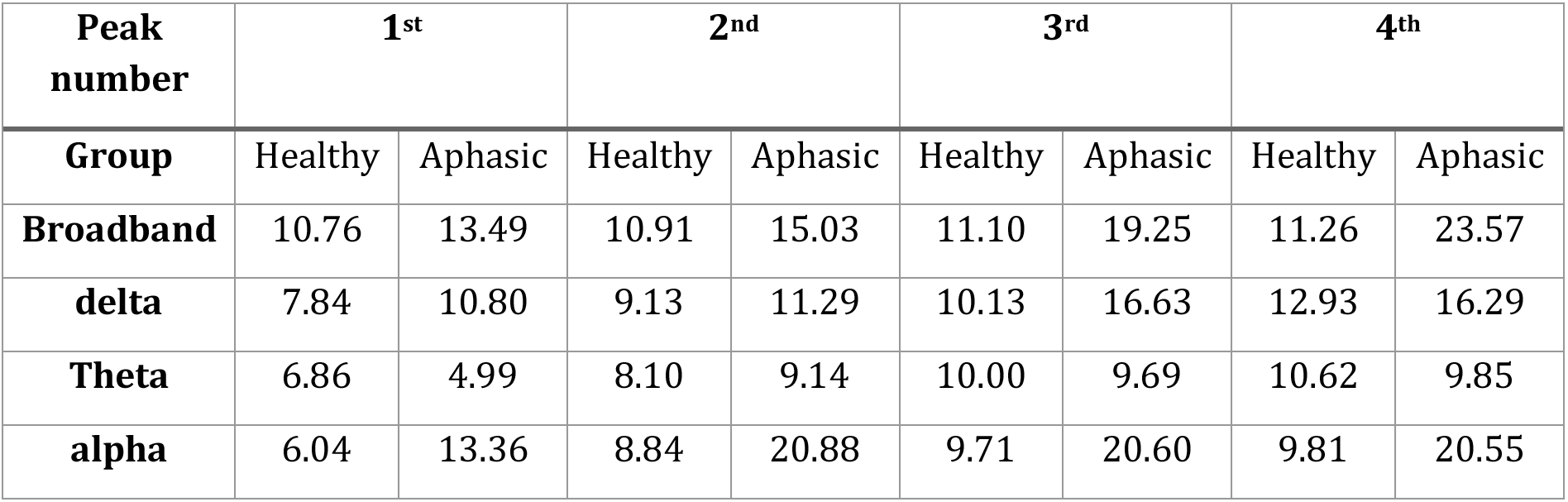
Consistency of the latency of the first four peaks of GFP patterns measured as the standard deviation of the peak’s latency for 10 random subsets of 25 epochs (in ms).

| **Peak number** | **1<sup>st</sup>** |  | **2<sup>nd</sup>** |  | **3<sup>rd</sup>** |  | **4<sup>th</sup>** |  |
| --- | --- | --- | --- | --- | --- | --- | --- | --- |
| **Group** | Healthy | Aphasic | Healthy | Aphasic | Healthy | Aphasic | Healthy | Aphasic |
| **Broadband** | 10.76 | 13.49 | 10.91 | 15.03 | 11.10 | 19.25 | 11.26 | 23.57 |
| **delta** | 7.84 | 10.80 | 9.13 | 11.29 | 10.13 | 16.63 | 12.93 | 16.29 |
| **Theta** | 6.86 | 4.99 | 8.10 | 9.14 | 10.00 | 9.69 | 10.62 | 9.85 |
| **alpha** | 6.04 | 13.36 | 8.84 | 20.88 | 9.71 | 20.60 | 9.81 | 20.55 |

This strong consistency in the latency of the peaks of GFP reflects an important synchrony of brain states across subjects and across trials. Hence the latency of GFP pattern (and the associated cortical microstates) is unlikely to predict the response latency of participants. To investigate the relationship between GFP curves and naming latencies, we compared the amplitudes of the peaks for short-latency trials (i.e., naming latency <1.5s) and long-latency trials (i.e., naming latency >1.5s). A two-way analysis of variance was carried out on the amplitude of the two main peaks of GFP (namely the second and third peaks post stimulus) with factors Latency (Short/Long), Group (Aphasic/Control). For both peaks, a significant main effect of Latency was found, for the third peak, a significant main effect of Groups was also found. No interactions between factors were observed (see table 3).

**Table 3.**
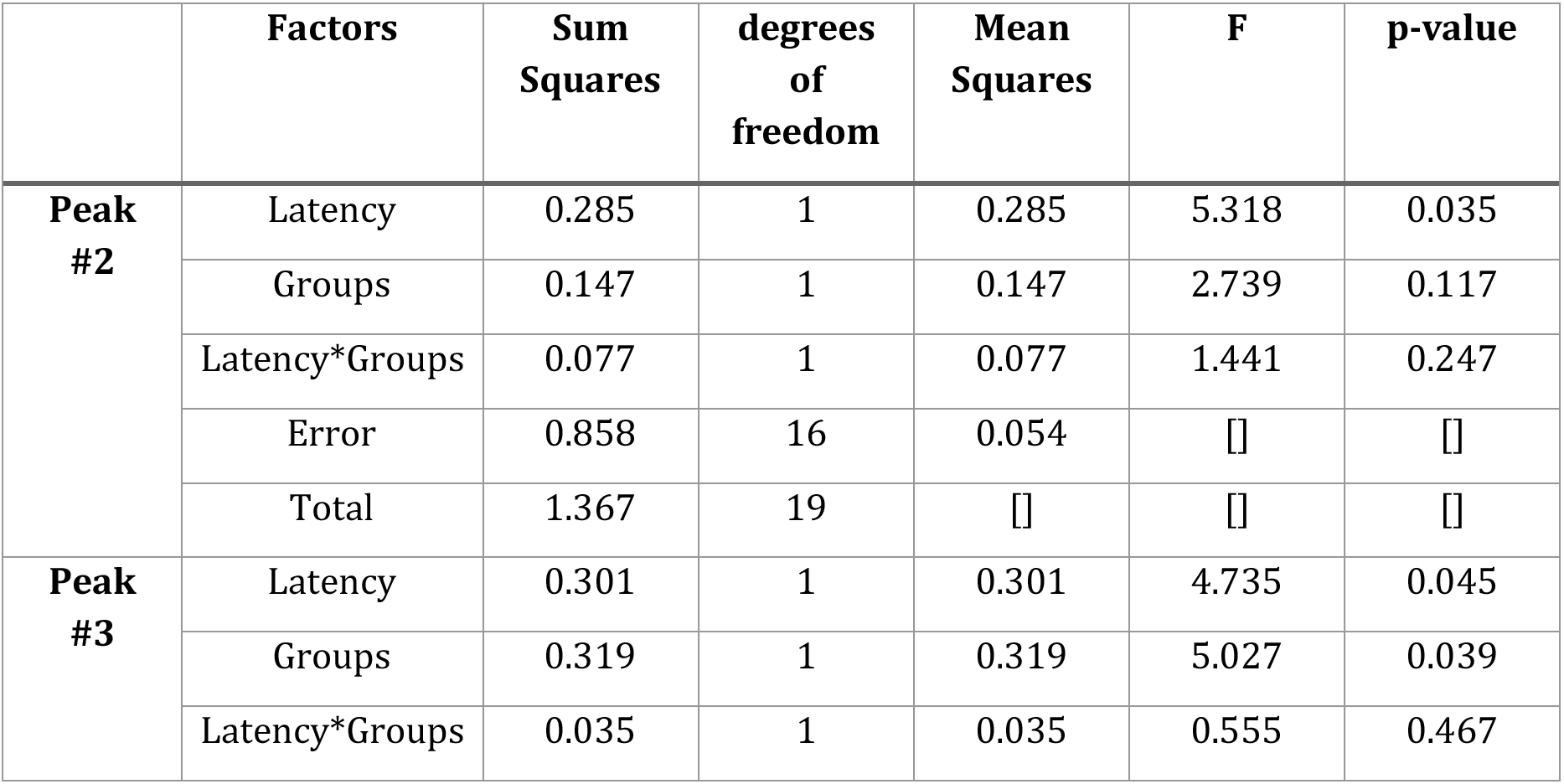
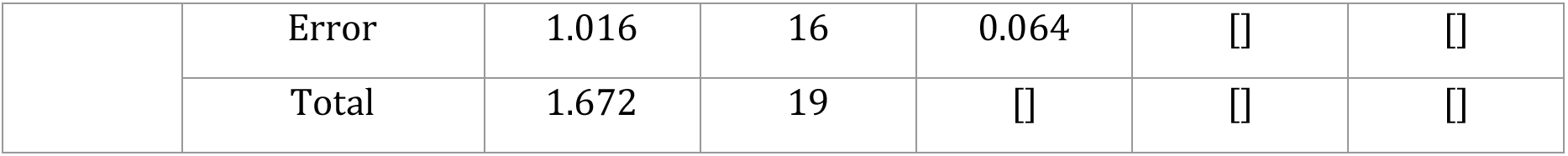
Output statistics from the two-way ANOVA performed across factors Groups, Latency for the two main peaks of the GFP pattern in the delta range.

|  | **Factors** | **Sum Squares** | **degrees of freedom** | **Mean Squares** | **F** | **p-value** |
| --- | --- | --- | --- | --- | --- | --- |
| **Peak #2** | Latency | 0.285 | 1 | 0.285 | 5.318 | 0.035 |
|  | Groups | 0.147 | 1 | 0.147 | 2.739 | 0.117 |
|  | Latency*Groups | 0.077 | 1 | 0.077 | 1.441 | 0.247 |
|  | Error | 0.858 | 16 | 0.054 | □ | □ |
|  | Total | 1.367 | 19 | □ | □ | □ |
| **Peak #3** | Latency | 0.301 | 1 | 0.301 | 4.735 | 0.045 |
|  | Groups | 0.319 | 1 | 0.319 | 5.027 | 0.039 |
|  | Latency*Groups | 0.035 | 1 | 0.035 | 0.555 | 0.467 |
|  | Error | 1.016 | 16 | 0.064 | □ | □ |
|  | Total | 1.672 | 19 | □ | □ | □ |

Post-hoc t-tests indicate that the amplitude of the second (average latency =290ms) and third peaks (average latency =424ms) of GFP in the delta range is significantly larger for the short-latency trials (Peak number 2, t(9)=2.43, sd=0.31, p=0.037; peak number 3, t(9)=2.89, sd=0.27, p=0.018). This may suggest that delta activity is involved in this specific cortical processing and that its contribution is larger for subjects with aphasia compared to control subjects. Even though the normality assumption was verified for this data, to account for the relatively low number of subjects, we also performed a non-parametric Kruskal-Wallis test to evaluate the effect of latency. It yielded a significant effect of latency for the second peak of GFP (χ²=4.17, df=1, p=0.0413) but not for the third (χ²=2.77, df=1, p=0.0903).

#### 3.3.2. Topographic representations

Figure 4 displays examples of ERP signals (top row) and the scalp maps computed at the peaks of the GFP (illustrated by vertical lines).

**Figure 4.**
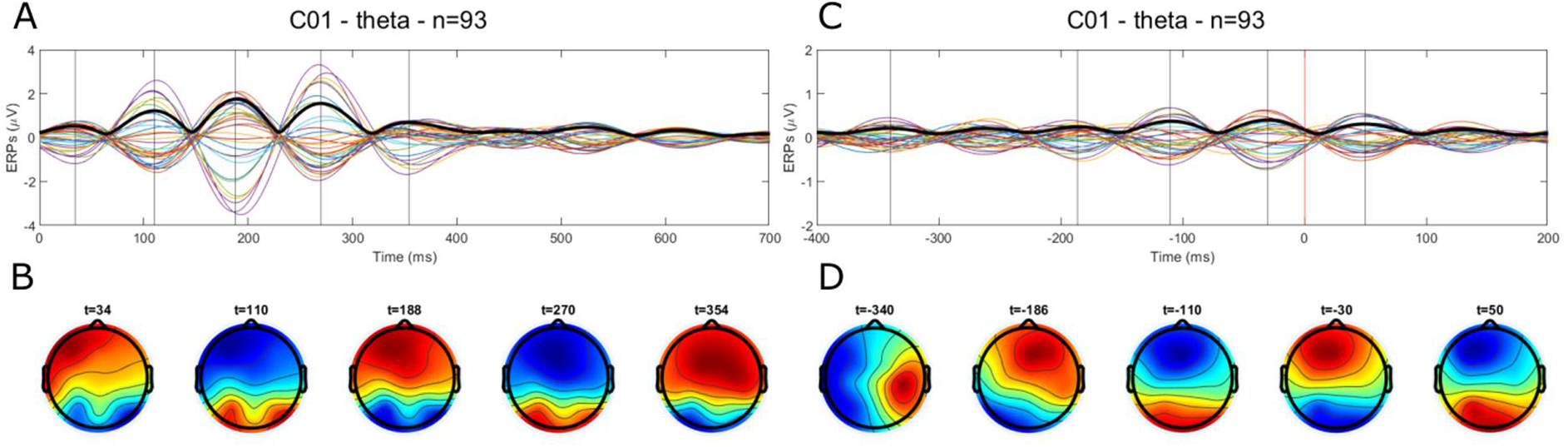
**(A)** ERP signals (colored lines) and global field power (thick line) for the stimulus-locked data. **(B)** Scalp maps corresponding to the timing represented by vertical lines in panel (A). **(C)** ERP signals (colored lines) and global field power (thick line) for the response-locked data. **(D)** Scalp maps corresponding to the timing represented by vertical lines in panel (C). Data for 93 successful trials of subject C01 filtered in the theta band.

Similar analyses were performed for all subjects and the same frequency ranges used in GFP analysis (see Section 2.4). In most cases, the same topographic organization was found in the delta, theta, and alpha ranges, with a dominant activity alternating between the occipital and the frontal area within the first 500ms after visual stimulus. This observation is consistent with the results reported by Laganaro, 2017 and Mheich et al., 2021 and is characteristic of the presence of oscillatory sources.

The response-locked patterns also revealed the activation of localized oscillatory sources. However, within the 200ms prior to the oral response these activations could mainly be observed in the frontal, parietal and central regions which also seem consistent with the source localization results from Laganaro, 2017. Interestingly, ERP signals could either show a strong synchrony across channels (i.e., in-phase channels) or a strong asynchrony (out of phase channels, see Figure S3).

### 3.4. Cortical dynamics

ERP measures provide an overview of potentially interesting patterns in the cortical activity. However, as argued by Alexander et al., 2013, averaging recordings such as cortical activity drastically reduces the intrinsic information. Since cortical activity is constantly changing and evolving, averaging multiple trials inevitably erases most of the information since it will only highlight the contribution of phase-synchronized physiological signals. In the following analysis, we aim at assessing the presence of regularity and repeatable patterns in the cortical dynamics at the single trial scale and the multiple trials scale using the amplitude trajectories obtained using the method described in Section 2.4.

Amplitude trajectories for each trial were projected on the parceled scalp map (Figure S4). Figure 5 displays six examples resulting from this method (C01, A01, A02, C04, A04, and A05). As an example, Figure 5.A shows that at t=252ms post-stimulus 45% of epochs showed a maximum of amplitude in the occipital region while for the remaining 55% of epochs, the peak amplitude was indistinctly distributed across the different scalp regions. At t=420ms 35% of epochs showed a peak amplitude in the central area and between 10% and 15% in the temporal and frontal areas.

**Figure 5.**
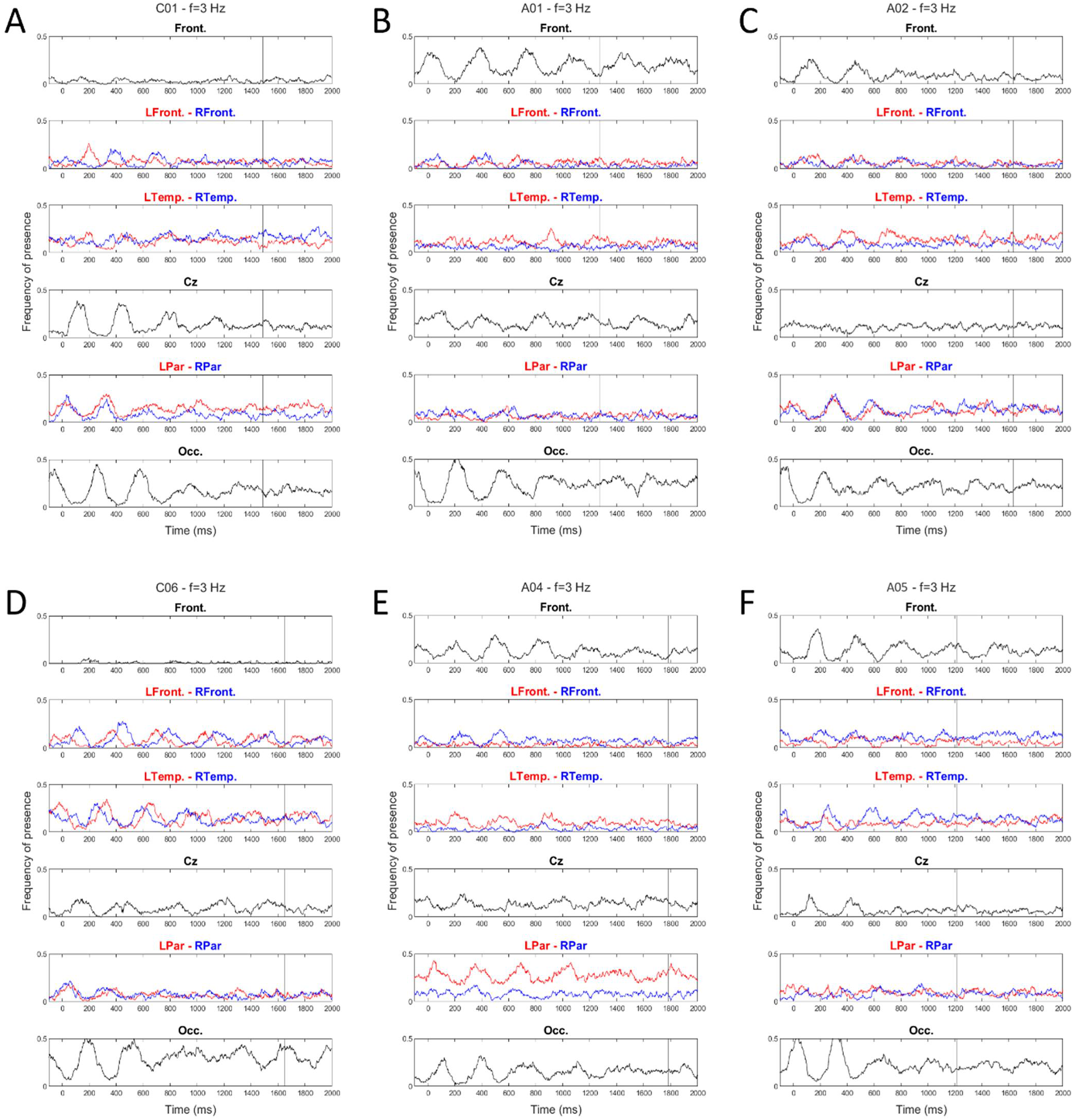
Spatial distribution of the maxima of amplitude. Frequency of detection of a peak of amplitude in different scalp regions as a function of time. Front= prefrontal area, LFront/RFront = Left/Right frontal areas, LTemp/RTemp=Left/Right Temporal area, Cz=Central area, LPar/RPar = Left/Right parietal areas, Occ.=Occipital area. Data from subject C01 **(A)**, A01 **(B)**, A02 **(C)**, C05 **(D)**, A04 **(E)** and A05 **(F)**, at a frequency of 3Hz and for all epochs, n=100.

The spatial distribution of the maxima of amplitude shows a strong across-trial regularity, especially in the occipital (“Occ.”) and central region (“Cz”), within the first 500ms which is coherent with the GFP patterns. It also indicates that cortical activity follows a clearly synchronized oscillatory pattern across trials. The noticeable delay between scalp regions also suggests a progressive propagation of cortical activity. Figure 5.A.B.C clearly reveal the alternating activity between the occipital and the frontal regions during the first 500ms post stimulus. This is consistent with the results from Salmelin et al., 1994 reporting a bilateral progression of cortical activity post-stimulus from the occipital region to the temporal and frontal regions. As in Salmelin et al., 1994, cortical dynamics during the overt speech preparation phases seemed much more variable across-subjects, and no clear pattern was observed during these phases.

All control subjects showed a weak contribution of the prefrontal area (e.g., Figure 5.A and D; “Front.”), while for all patients but A03, the presence of activity in the prefrontal area was observed (e.g., Figure 5 B, C, E, F). For all patients but A03 we can observe an oscillatory pattern of the activity between the prefrontal area and the occipital area, indicating a strong across-trial synchrony (e.g., Figure 5 B, C, E, and F). This observation was particularly obvious at a frequency of 3Hz (i.e., delta range) and could persist for the entire epoch duration. It may therefore explain the atypical pattern of GFP in the delta range for most patients mentioned in Section 3.3 as a potential signature of pathological activity. Similar analysis at higher frequencies also shows such oscillatory patterns but with a smaller amplitude. This results from the fact that trajectories at higher frequencies propagate faster which makes this measure more sensitive at high frequency. Moreover, a similar oscillatory pattern can be observed in control subject and A03 between central area and occipital area. Finally, the onset of the oscillating pattern could be observed during the 100ms prior to visual stimulus which may reflect an anticipation process.

Figure 6 displays six examples of unrolled trajectories to assess the presence of repeated patterns at the single-trial scale. Despite an important variability across subjects and across trials, several observations can be made from this representation.

**Figure 6.**
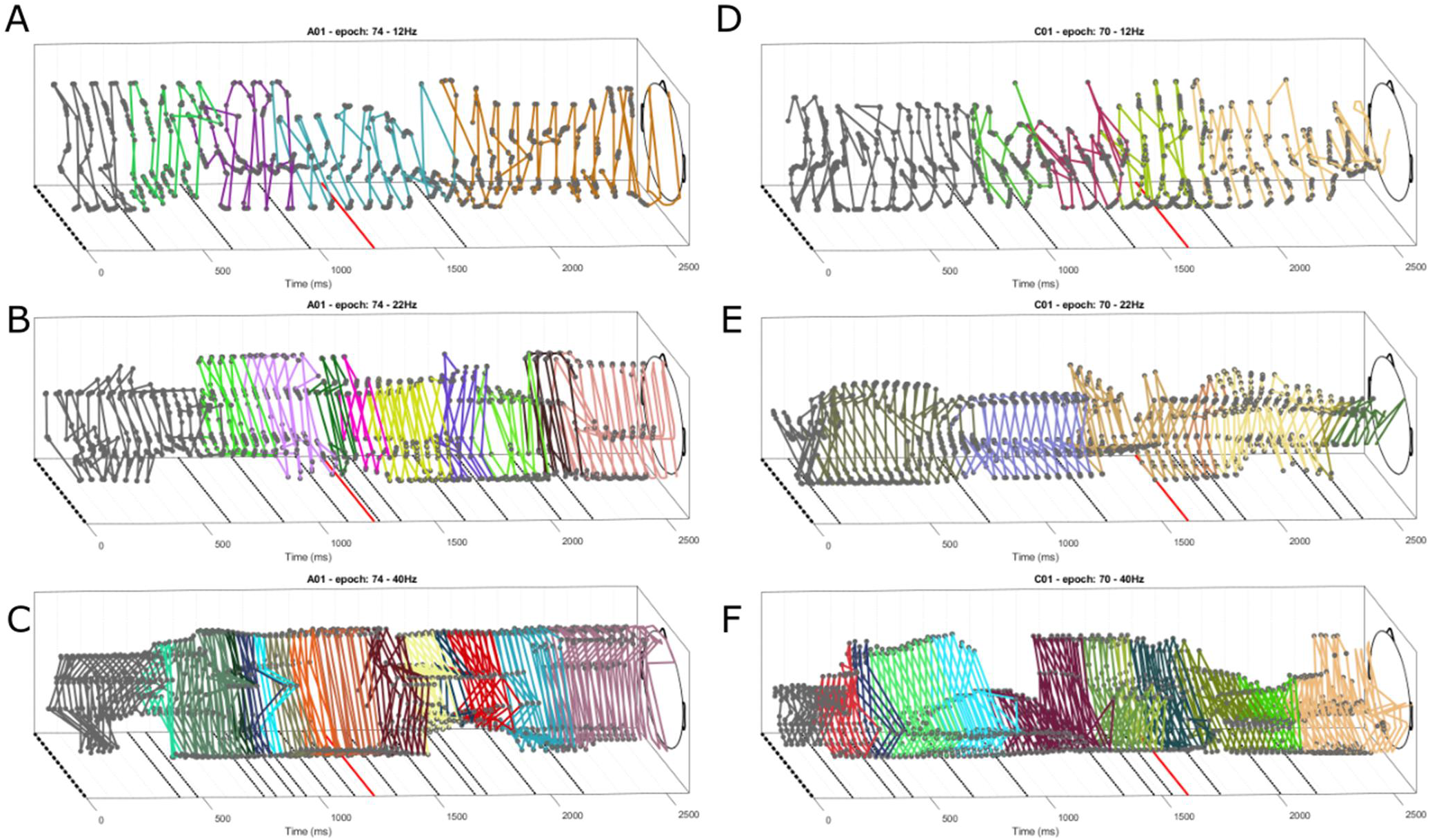
Unrolled trajectories for subject A01 **(A, B, C)** and C01 **(D, E, F)**, for the same word “chien” (i.e., dog), at 12, 22, and 40 Hz from top to bottom. In each frame, the dashed line at t=0ms represents the image presentation, the red line represents the voice onset, and dotted lines represent the abrupt changes in trajectory detected using the Hausdorff distance measure. Different colors of the trajectory curve represent the corresponding segments. Cartoon head is shown for visualization clarity.

First, the trajectories cover and connect different regions of the scalp. These connections may be sudden and at a large scale (e.g., frontal-occipital) or progressive and at a smaller scale (e.g., successive samples in a restricted regions). Second, trajectories form specific patterns of activity which are periodically repeated with a strong regularity. Each of these patterns seems to connect a limited number of brain regions. This periodic organization is consistent with the presence of an oscillatory activity, for the duration of each pattern (referred to as a cycle in the following) equals Tcycle = 1/f. Each pattern consists of the maximum of amplitude traveling along a particular pathway. The repetition of a specific pattern for several cycles is referred to as a segment. The durations of these segments was highly variable and ranged from tens of milliseconds to several hundreds of milliseconds. Longer segments could thus correspond to relatively demanding cognitive tasks being executed while shorter ones are more likely to reflect switches in the global cortical processing. Third, the fact that most patterns form approximately closed polygons over one cycle relate to the relationship between frequency and wave velocity, in a particular manner that the oscillation period covaries with the spatial period. In other words, an oscillatory activity may propagate along a specific pathway whose length is dependent on the oscillation frequency. This observation is in agreement with Ermentrout & Kleinfeld, 2001b and seems also verified in the data shown in the literature (Alexander et al., 2013; Zhang et al., 2018; Alamia et al., 2019).

We performed the automatic segmentation of the unrolled trajectories presented in Section 2.4.3 which can be seen as a single-trial cortical microstate as opposed to the microstates identified from ERP analysis (Hassan et al., 2015; Mheich et al., 2015; Laganaro, 2017). Dotted lines along the times axis in Figure 6 represent these boundaries and different colors illustrate the different segments.

As a matter of comparison, we performed the segmentation method across the entire experiment (100 trials), for each subject and frequency. We then counted the number of segments from the image presentation to the voice onset. It results that the median of the distribution of number of segments for each subject varied from 4 to 7, which is also coherent with the observation of Mheich et al., 2021 using a different paradigm (phase locking value) and considering a different frequency range (30-45Hz).

Figure 7 represents the distribution of the number of segments boundaries for all subjects and all trials (n=10*100), considering 50ms bins. Overall, this distribution could be described as a combination of a baseline trend and a periodic component. The periodic component can be explained by the fact that the duration of segments is dependent only on the oscillation frequency, and by the relatively strong inter-subject and inter-epoch synchrony of the cortical activity. It is also worth noting that this segmentation method revealed several frequency-dependent features. First, the segmentation was more obvious for high frequencies (22, 30, 40Hz) than for lower frequencies. More precisely, the estimated duration of each segment decreased with the oscillation frequency and could be fitted with a quadratic law (*tsegments* = 0.5292*f^2^ - 38.78*f + 830.5, r²=0.998). Second, we can observe different properties of distributions of segment boundaries for different frequencies (Figure 7). At 3Hz, the distribution peaked around 500ms and 1500ms which might relate to the end of the initial recognition processing and to the speech production respectively. At 9Hz, the periodicity of the distribution was much less consistent around the median naming latency (i.e., between 1100ms and 1550ms). This might reflect inter-trial and inter-subjects’ variability in the voice onset time. For frequencies of 9, 12, 16 and 22Hz the baseline trend gradually increased from the image presentation (first vertical line, t=0ms) and plateaued around the median latency (second vertical line, t=1456ms). This behavior was particularly obvious at 16Hz and 22Hz and indicates a strong relationship between the cortical dynamics and the task. In contrast, at 30Hz and 40Hz, an obvious increase in boundaries detection was seen within the first 100ms post-stimulus likely due to an early neural response to the visual stimulus while from 200ms onwards, the baseline remained constant. This suggest that the cortical activity at higher frequencies (30 and 40Hz) is less dependent on the task being executed.

**Figure 7.**
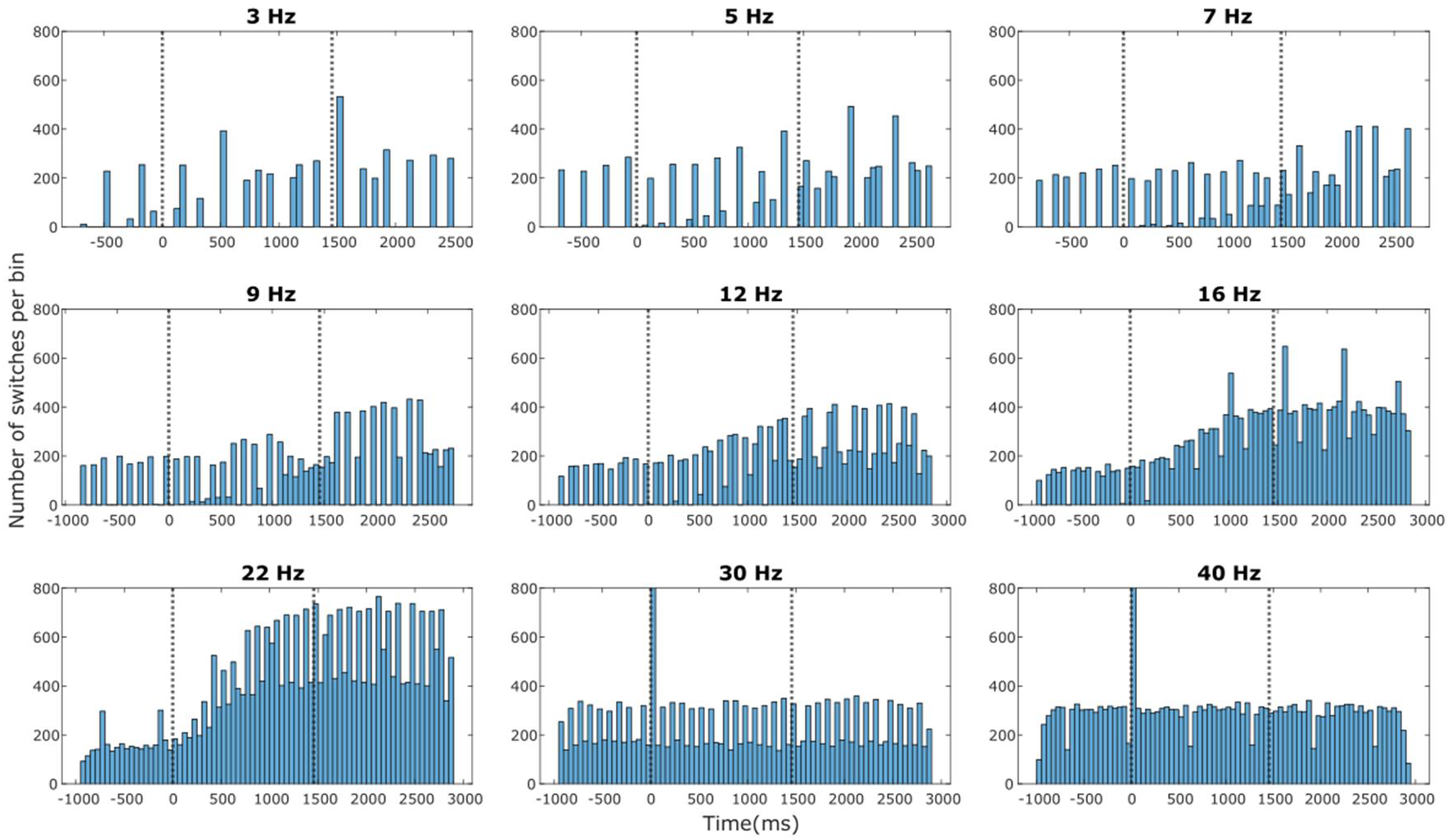
Histograms of the number of segment boundaries calculated for all ten subjects, all 100 epochs and for all nine frequencies (3, 5, 7, 9, 12, 16, 22, 30, and 40Hz). Bin width is equal to 50ms. Vertical lines indicate the image presentation (t=0ms) and the median latency (t=1456ms) respectively. For 30 and 40Hz, the histogram at t=0 exceeded 800 but the same scale was used for clarity.

### 3.5. Detection of traveling waves (TW)

In the following, we present the results from the analysis methods of the cortical dynamics used to detect and characterize TW based on the amplitude and the instantaneous phase of the EEG signals. Figure 8.A displays one example of an amplitude wave captured using the method described in Section 2.4. As a matter of validation, we compared this amplitude-based method with a phase-based approach.

**Figure 8.**
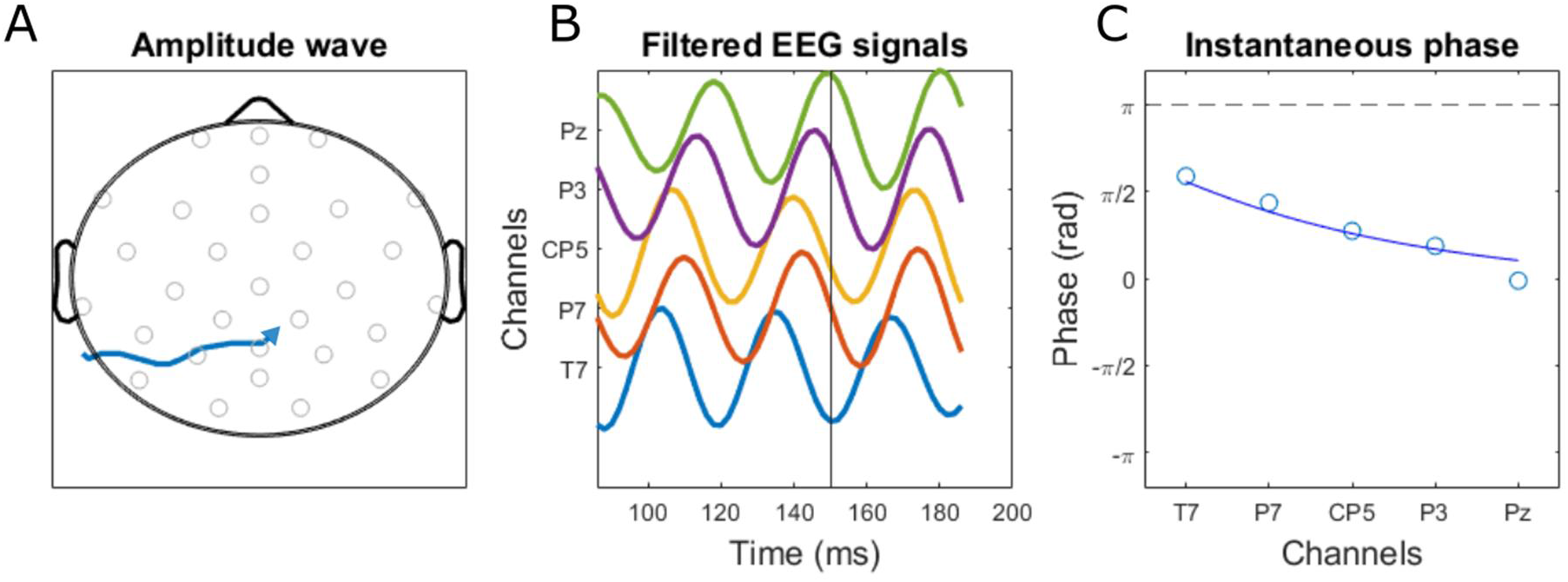
**(A)** Topographic representation of one amplitude wave (blue arrow). Grey circles indicate the position of EEG electrodes. **(B)** EEG signals from channels T7, P7, CP5, P3 and Pz during the same time window. **(C)** Instantaneous phase of channels T7, P7, CP5, P3 and Pz (circles) and results from the circular-linear regression (line). Data recorded with subject C01 and filtered at 22Hz.

Overall, for all frequencies used in the present study 58% of the amplitude waves yielded a regression coefficient higher than 0.8. Figure 8.B displays the EEG signals measured at electrodes T7, P7, CP5, P3 and Pz (i.e., along the trajectory of the amplitude wave). Figure 8.C displays the instantaneous phase on these same channels and the result from the circular-linear regression model. The present method enables to quantify the number of TW, approximate their length and their velocity. The length and velocity were estimated by considering the scalp linear distance (Iz-Fpz) equal to 30cm. The propagation velocity as a function of frequency could be fitted with a linear curve which is consistent with several previous results (equation 1).

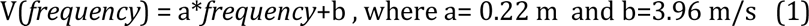

Equation (1) is consistent with several previous results (Figure 9.A, Patten et al., 2012; Muller et al., 2018b; Zhang et al., 2018). Quantitatively the estimated velocity ranged from 1 to 15 m/s with an average value of 5.97m/s for a frequency of 9Hz which is also coherent with the results from Patten et al., 2012 (6.5m/s for alpha waves), Massimini et al., 2004 (1.2–7.0 m/s for slow waves), Botella-Soler et al., 2012 (1m/s for delta waves, note that the authors argued that the wave velocity was probably underestimated) and consistent with the observation of macroscopic waves (Muller et al., 2018a). The length of the amplitude waves (Figure 9.B) was also frequency-dependent with faster waves propagating over longer distances.

**Figure 9.**
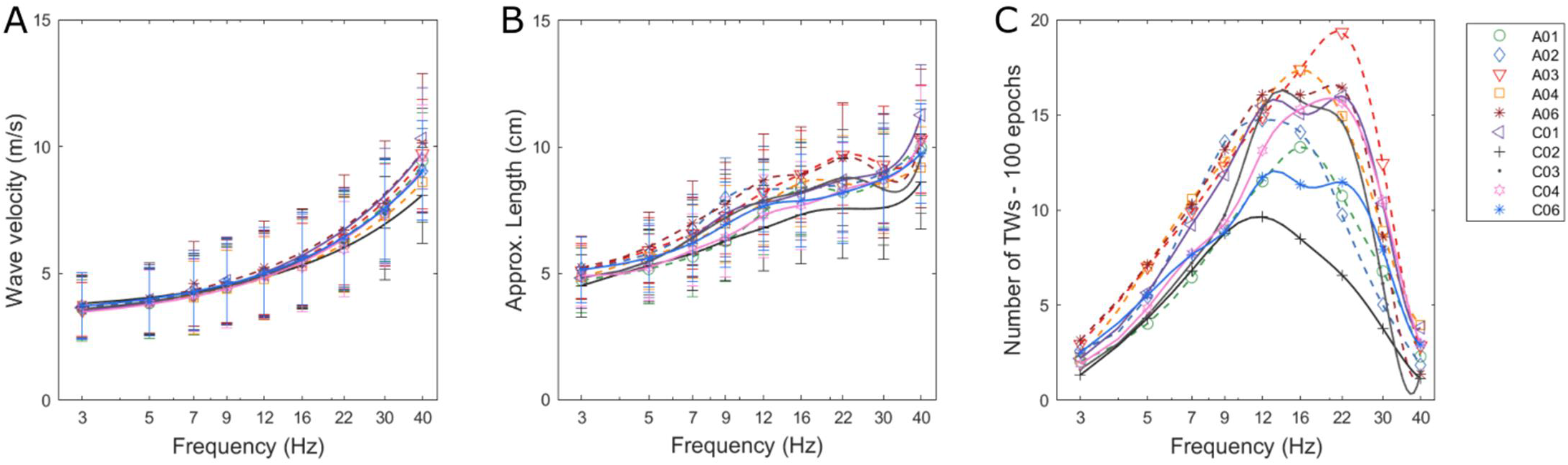
**(A)** Approximate velocity (in m/s) of the TW as a function of frequency. **(B)** Approximate length (in cm) of the TW as a function of frequency. **(C)** Normalized number of TW as a function of frequency (in /s). Different symbols indicate different subjects. Data from subjects with aphasia are represented with dashed lines. n=100 for each condition.

Figure 9.C displays the number of TW as a function of frequency for all subjects. It is worth noting that this number did not peak at a frequency corresponding or close to the alpha peak frequency, meaning that high cortical activity is not necessarily associated with a larger number of TWs. All subjects exhibited a maximum of waves between 12 and 30Hz. Interestingly, six subjects showed a typical profile exhibiting a peak around 22 Hz while the other four subjects (A01, A02, A04, and C02) showed either a plateau or a roughly gaussian distribution centered between 12 and 22 Hz. Note that a very similar velocity profile was found from the analysis of resting state data (eyes-open). It was also found that the number of detected waves, normalized by the duration of considered dataset (i.e., expressed in number of TWs per sec) did not change between the resting state and the naming task conditions.

### 3.6. Mapping cortical activity

Our results indicate that cortical TW can be detected from EEG signals. Finally, we aimed at clarifying the role of TW within the cortical network. To investigate cortical connectivity, we performed the detection of TW using the algorithm described above for each subject, at each frequency and for both the naming task and the resting state condition (eyes-open). Then, detected TW were plotted as a 2D scalp map. Figures 10.A and B display the TWs identified in 10 occurrences of the control word “chien” (i.e., “dog”) from t=0 to t=2500ms at a frequency of 22Hz for subjects A01 and C01 respectively. The color scale illustrates the timing of the different waves, relative to the end of the TW. Figure 10.C, D represent the same analysis for three minutes of resting-state data. Both resting state (figure 10.C, D) and naming task data (Figure 10.A, B) showed that TW could be detected across the entire scalp as well as an important inter-subject variability of the cortical dynamics (Massimini et al., 2004; Murphy et al., 2009; Botella-Soler et al., 2012; Alexander et al., 2013; Salmelin et al., 1994). It is worth noting that for lower frequencies, (5, 7 and 9Hz) TW were more spread across the entire scalp, while at higher frequencies (22 and 30 Hz) more obvious and redundant connections could be seen.

**Figure 10.**
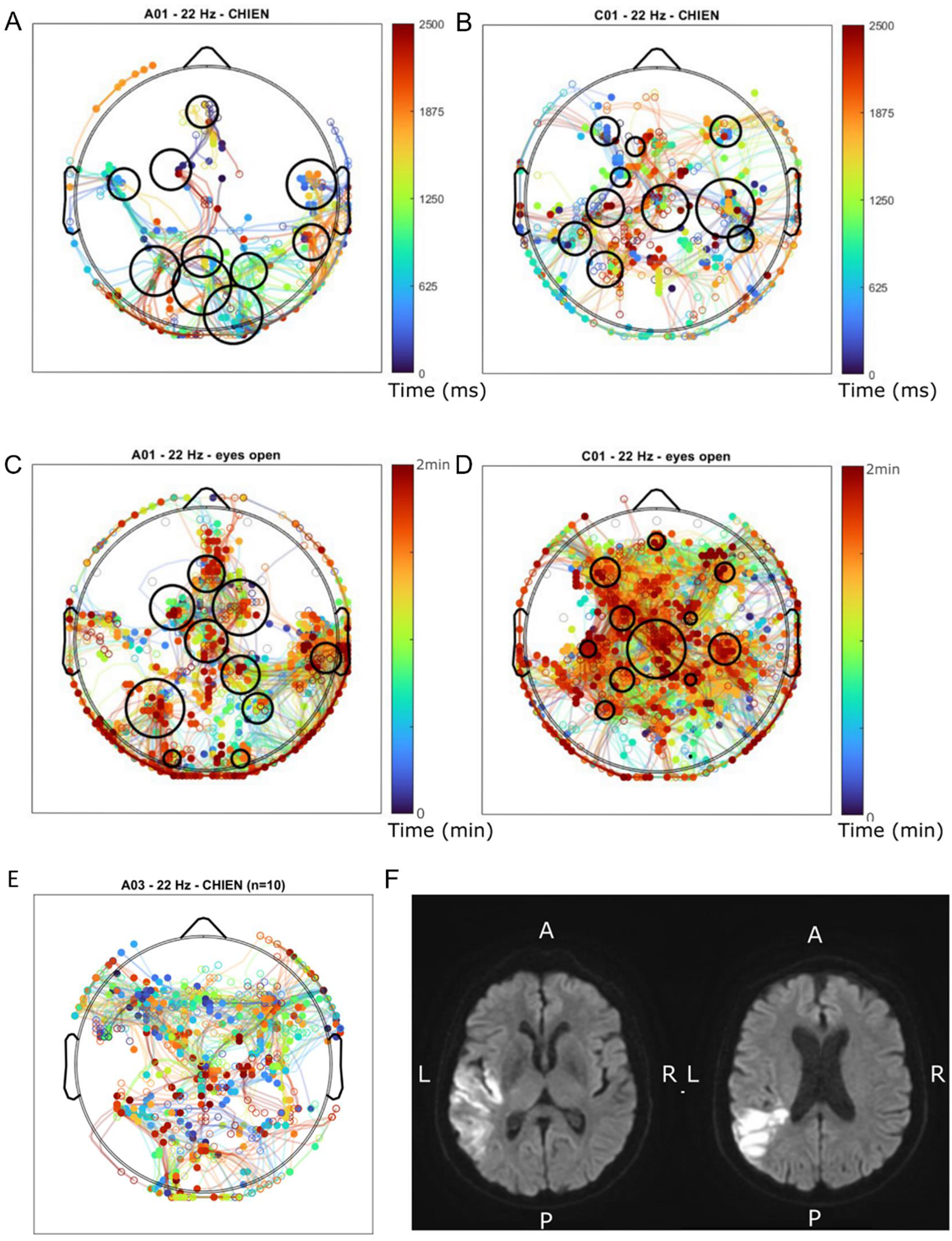
(A), (B) TW detected in 10 repetitions of the word “chien” (i.e., dog) for subject A01 and C01 respectively, at a frequency of 22Hz. Colormap indicates the timing of the end of the wave relative to the visual stimulus. Black circles illustrate main hubs identified from density maps analysis. **(C), (D)** TW detected during two minutes of resting state recording with eyes open for subject A01 and C01 respectively, at a frequency of 22Hz. **(E)** TW detected in 10 repetitions of the word “chien” (i.e., dog) for subject A03. **(F)** MRI images from subject A03 showing the location of the stroke lesions in left temporo-parietal region.

In contrast, the distribution of TW showed a remarkable within-subject consistency. Our analysis also revealed the presence of regions where many TWs can be detected (black circles in figure 10.A). Circles in figure 10 result from the hubs detection procedure described in Section 2.4 and the circles radii are proportional to the sample size in this bin. If TWs play a role in the communication and synchronization between key areas, such crossroads may thus represent relevant hubs. The locations of main hubs were very consistent across frequencies up to 22Hz. The largest deviation in the hub locations was found for a frequency of 30Hz. In addition, the distribution of these hubs was much more homogeneous for healthy subjects than for subjects with aphasia (see examples at 22Hz for all subjects in Figure S5).

For all participants with aphasia, an asymmetrical or inhomogeneous distribution of TW and their associated hubs could be noted (e.g., subject A01 in Fig.10). We assessed the correlation of this observation with the position of the stroke lesions by analyzing Magnetic Resonance Imaging (MRI) images and clinical reports. Overall, this analysis showed a sparser density of TW (missing hubs) in the lesion area sometimes associated with a higher activity in the perilesional areas (displaced hubs). More specifically, A03 and A05 mainly showed a sparser density of TW in the vicinity of their stroke lesions (i.e., left temporal and parietal areas for A03, left-temporofrontal area for A05). A01, A02 and A04 showed an important lack of TW both in the lesion area as well as in the homologue region. This seemed to be compensated by a higher activity in either the perilesional area or the central area. This reorganization of cortical activity also echoes with the observation made in section 3.4 stating that for most patients with aphasia the amplitude trajectories covered the anterofrontal region which is rarely seen in our group of control subjects (see also Salmelin et al., 1994). Figure 10.E.F shows the distribution of detected TW for subject A03 and the MRI images showing the location of the stroke lesions.

The present methods and results are reminiscent of the study by Salmelin et al., 1994 using magnetoencephalography. We also compared the locations of these hubs with maps of the human connectome reported by Kabbara et al., 2017 and van den Heuvel & Sporns, 2011. This comparison exhibits a striking overlap between the hubs identified by TW and either the resting state network, the default mode network, or the rich-club network (see Figure 7.B in Kabbara et al., 2017). However, it is known that these distinct networks may partially overlap and change over time. Hence it remains complex to determine to which network our dynamic network belongs to.

## 4. Discussion

In this study, we were interested in investigating the spatiotemporal dynamics of cortical activity during language production in the healthy and lesioned brain. Healthy volunteers and people with post-stroke aphasia performed a self-paced picture naming task during EEG. Our main findings were as follows: First, we observed strong synchronization of cortical oscillations in the first 600ms post-stimulus, with a time shift between both groups. ERPs and corresponding brain microstates indicate coordinated brain activity alternating mainly between frontal and occipital zones resulting in standing waves. Second, TW were identified from both phase and amplitude analyses at the single trial level. The spatial distribution of TW was altered for participants with aphasia. Third, the presence of TW in different cortical areas showed a remarkable time coordination, essentially between frontal and occipital zones, and in relation with GFP oscillations. The spatial dynamics differed between participants with aphasia and control subjects, with prefrontal TW being selectively present in participants with aphasia. Collectively, our results show that TW contribute to the synchronization and communication between different brain regions by interconnecting cortical hubs. Moreover, our findings imply that such dynamics are affected by brain lesions.

### 4.1. Behavioral correlates of language production

Behavioral results indicate that participants with aphasia could perform well despite showing signs of effortful processing. Most naming errors could be categorized as semantic or phonemic paraphasia. A significant difference in naming latency was observed between healthy participants and participants with aphasia for all trials as well as for successful trials only. Note that the naming latencies measured in the present experiment are longer than those reported in the literature (approximately 800ms across five different studies reviewed by Laganaro, 2017, and 883ms on average for Alario & Ferrand, 1999). The main reason that could explain this difference is that subjects in both studies were younger than in the present study (25 years old on average across the five studies reviewed in Laganaro 2017, 46 university students in Alario et al., and 55 years old on average in the present study). An alternative explanation for the observed longer latencies in our study may come from task design which allowed self-paced production without emphasizing the speed of the response, indeed the participants were not asked to answer as fast as possible. To account for longer latencies Roelofs et al., 2017 proposed the possibility to scale the model of Indefrey and Levelt. However, such transposition is complex since the origin of this delay is unclear. Besides, ERP analysis suggests that the timing of the initial cortical processing was relatively consistent with previous studies.

### 4.2. Electrophysiological correlates of language production

EEG data were analyzed using a static and a dynamic approach to improve our understanding of cortical dynamics during language production. While the functional role of alpha oscillations (8-12 Hz) is still unclear (Klimesch et al., 2007; Klimesch, 2018; Alamia et al., 2019), the dominant alpha peak frequency is often used as a reference in EEG studies since it is consistently present in most human subjects. Petrovic et al., 2017 reported that post-stroke subjects, in the sub-acute phase show lower peak-frequencies than healthy subjects. Although our analysis of peak frequency allowed detecting a lower group mean of peak frequencies for the chronic-stage participants with aphasia, our small sample size did not yield any significant group effect.

ERP analysis enabled the replication of several results from the literature despite a limited cohort of participants and a relatively low-resolution EEG system (i.e., 32 electrodes). Cortical microstates can be identified during the cortical processing of picture naming. In the low-frequency bands (delta, theta), the main peaks of the GFP were associated with a marked alternating pattern between the occipital cortex and either the anterior frontal or the dorsolateral frontal cortex. From 600ms onward the intersubject variability was much larger thus limiting the possibility to draw global conclusions (Salmelin et al., 1994; Alexander et al., 2013, Saravani et al., 2019).

The present results demonstrate the strong within and between-subjects consistency in the latency of the GFP patterns within the first 600ms post-stimulus. They also show some consistency with previous studies confirming the important regularity in post-stimulus cortical activity. A recent meta-analysis on five different picture naming studies by Laganaro, 2017 reported that, at the group level (n=118), the average (broadband) GFP pattern exhibited a very consistent peak at approximately 110ms post stimulus and less salient peaks centered at approximately 225ms and 340ms post stimulus respectively. In Mheich et al., 2021 (23 participants naming 148 objects) the GFP exhibited a dominant peak at 180ms as well as multiple peaks centered around 110ms, 130ms, 210ms and 230ms post stimulus. Due to a high level of consistency between subjects with aphasia and control subjects, the GFP latency itself is unlikely to predict difficulty or failure in the picture naming cortical processing.

According to Indefrey and Levelt 2004 and Laganaro, 2017, the first stage of the picture naming process relates to the image and concept recognition, the second peak of GFP occurring around 200ms may correspond to the lexical selection process while the following segments (third, fourth peak) may relate to the phonological and phonetic encoding and monitoring (Indefrey, 2011). The potential need of rescaling this model to account for longer naming latencies could be considered here.

The analysis of ERP and GFP in limited frequency ranges provided additional information. We found that trials with a shorter naming latency were associated with a higher amplitude of the second and third peaks of the GFP pattern in the delta range (between 290ms and 424ms post stimulus). Despite some discrepancies in naming latency, these time windows may relate to semantic (“Lexical phonological code retrieval”) and phonological (“syllabification”) processes (Indefrey et al., 2004; Llorens et al., 2011; Indefrey, 2011; Croce et al., 2020). This observation is consistent with the fact that in the present experiment, most naming errors resulted from semantic paraphasia (i.e., correct semantic context but inaccurate target selection) and/or phonemic paraphasia (substitution or rearrangement of phonemes). This increase in amplitude was even larger for participants with aphasia which might be the signature of a pathological slow activity. This observation was corroborated by the fact that, overall, the GFP patterns for participants with aphasia showed a larger and/or delayed contribution in the delta range.

### 4.3. Amplitude trajectories allow spatiotemporal tracking of language production

The amplitude tracking method used here enabled to further investigate the distribution of cortical activity and its across-trial consistency with more accuracy than the ERP approach.

Figure 5 reveals a very consistent oscillatory pattern between the occipital and both central and frontal area for control subjects. This might indicate the early involvement of premotor areas (Llorens et al., 2011). In contrast, participants with aphasia (except A03) showed enhanced delta activity in the prefrontal cortex. As an important caveat, this observation does not mean that the prefrontal cortex is not active in control subjects but rather that the cortical activity was consistently larger in amplitude in the central and frontal regions.

The illustration of amplitude trajectories (Fig. 6) indicates the presence of stable spatiotemporal segments of the cortical process lasting up to several hundreds of milliseconds. These patterns are consistent with an oscillatory activity propagating through a spatiotemporally coherent pathway. Overall, the distribution of the segment boundaries in time (Fig. 7) provided frequency-specific markers which could be related to the various steps of the picture naming task, even though further investigation is required to strengthen our findings.

### 4.4. Traveling waves allow the identification of hub regions

Applying restrictive criteria on the amplitude trajectories enables to identify propagating TW in two dimensions. The estimated velocity, length and number of TW are consistent with previous results from the literature (Zhang et al., 2018). We also combined amplitude tracking and phase regression methods to strengthen our observation. Results show that in around 60% of cases, the detection of an amplitude wave was also coherent with the detection of a phase wave. The residual 40% of amplitude waves which could not by associated with a phase wave can be explained by the fact that the two methods are intrinsically different. Indeed, amplitude waves are defined by concatenating successive samples. On the contrary, phase waves are identified when the instantaneous phase of several successive electrodes forms a regression. For example, if the maximum is not moving, there is no amplitude wave, but there may be a phase wave. Conversely, the phases of the successive electrodes may be the same (no phase wave), but the maximum amplitude may shift. Besides, it is likely that other dynamic patterns or propagating waves occur at lower amplitudes. These patterns would therefore not be captured by the present technique.

The distribution of TW demonstrates a strong activity on the entire scalp surface. We also noted the existence of limited regions with a higher density of TW. The location of these hubs demonstrated a strong within-subjects consistency for different frequencies both at rest and during the picture naming task. Their number and position is also consistent with either connector hubs of the default-mode network, resting-state network and/or rich-club network (van den Heuvel et al., 2011; Sporns, 2013; Kabbara et al., 2017).

The present results provide evidence suggesting that TW detected from amplitude tracking ensure the connection between at least main connector hubs of these resting state networks. In particular, the frequency dependence of TW length and their spatial distribution could indicate that provincial hubs are linked by slow TW while connector-hubs are connected by faster (higher-frequency) waves.

Data in participants with aphasia showed specific changes in the distribution of TW in the lesional and perilesional area. This supports the fact that the properties of damaged neural tissues may be modified by the stroke and further neural degeneration (Bessonov et al., 2019). The presence or absence of TW in the lesional area may reflect different degrees of initial tissue damage and potentially different degrees of neural self-regeneration (Lindvall et al., 2015). Again, the absence of TW detected with the present method may also indicate that the cortical activity in these regions is lower in amplitude and thus not captured by our technique.

Our data further suggest that, in response to this relative lack of TW in the vicinity of the lesioned tissue, local increase of traveling waves are observed in homologous regions of perilesional regions.

This may also indicate that stroke lesions can affect cortical hubs and may thus induce the displacement of one or several main hubs, or even their suppression. This obviously questions the consequences of such drastic changes in the cortical organization.

However, one must bear in mind that while the approximate location of stroke lesions could reliably be associated with a sparser distribution of TW, some “blank” regions could also be seen in other regions and/or in healthy subjects. This observation thus suggests that the distribution of TW highlights the presence of main pathways across the scalp resulting from years of cortical plasticity which is even more pronounced in the presence of stroke lesions. Reproducing this analysis of cortical dynamics longitudinally from the acute to the chronic phase could provide valuable insights for our understanding of the cortical post-stroke reorganization.

Our observations suggest that dynamic hubs identified throughout this study act as major oscillatory sources of cortical activity. Therefore, they seem to initiate and receive most of the (high-amplitude) TW. This finding strengthens the hypothesis that cortical TW contribute to the synchronization of brain areas and suggests that part of their role consists in connecting hubs of the default mode network.

However, at this stage the specific role of these hubs and neighboring regions is difficult to determine. It is still unclear whether the sequential order of activation of these hubs is important for efficient cortical processing. Likewise, our results do not allow to differentiate between serial and interactive theories of picture naming. We note that our study was not designed to provide evidence for or against a specific model. Rather, we aimed to probe the cortical dynamics of distributed processing in healthy and lesioned brains during a prototypical language production task. A hierarchal activation of hubs could not be established, nor to clearly demonstrate how TW relate to picture naming processing for multiple reasons. First, as reported by numerous previous studies cortical dynamics show an important inter- and intra-subject variability (e.g., Alexander et al., 2013; Croce et al., 2020; Saha et al., 2020; Salmelin et al., 1994). Second, the present approach does not account for frequency coupling or phase-amplitude coupling mechanisms which may play an important role. The movie presented in supplementary materials (Movie S6) illustrates how TW at different frequencies may merge in a given hub or how a TW at a given frequency can pursue its propagation at a slightly different frequency. Such complex mechanisms should be taken into account, and further studies are needed to deepen our understanding of cortical dynamics.

### 4.5. Limitations

Our results are limited by the small sample size and the relatively large number of outcome variables. Consequently, our findings should be interpreted with caution. We further note that, in the present study, EEG was recorded with a relatively low spatial resolution (30 active contacts, while HD-EEG can use as much as 256 contacts). EEG data were thus interpolated from the 30 contacts. Besides, the projection of the EEG data on the scalp is not accurate enough to enable identifying the corresponding anatomical regions. In particular, the topographic maps, trajectories, and TW at the edges of the interpolation grid must be carefully interpreted since the accuracy of the interpolated data cannot be predicted. It is therefore possible that using a higher density of electrodes could provide finer details and facilitate the interpretation of such results.

### 4.6. Conclusions

Overall, the present observations support the fact that the initial steps of the picture naming cortical processing are operated with a strong time regularity. Our results further indicate that cortical activity follows periodic patterns of activation (referred to as segments) during periods of time which may last from 700ms to 100ms as the frequency increases from 3 to 40Hz. While some similarities could be found between trials, these segments are highly variable.

The cortical dynamics within each segment could be described as a combination of relatively stable brain sources (recurrent hotspots in the TW topographic maps) potentially resulting from the synchronous and periodic firing of a local neural population inducing standing waves (Bhattacharya et al., 2022). In parallel, continuous connections could be observed across the scalp which may indicate the existence of cortical TW. As a result, the peaks of ERP and GFP patterns mainly picture the influence of such sources.

Additionally, we found that the distribution of changes in amplitude trajectories (i.e., segments boundaries) followed a periodic pattern related to the segments’ length. Such a behavior suggests that each processing step is synchronized and limited in time. It therefore seems that the role of TW is to ensure the synchronization and involvement of relevant cortical areas (hubs). Each segment of the unfolded trajectories seems to correspond to multiple iterations of a specific operation. Once a response threshold is reached, a new segment appears. Then multiple iterations of a new operation begins. The production of a correct or false response will then depend on the efficacy of the processing (memory retrieval, inhibition of incorrect objects). Collectively, the present results improve our understanding of cortical dynamics as well as the influence of stroke lesions on oscillatory activity. Our results and methods provide some useful hints for further investigation. Replicating such analyses for a semantic or phonological task, and over a larger cohort of participants, could highly improve our understanding of the task-specific cortical processing.

Our results may also have direct implications for future modeling studies on the cortical dynamics and the role of TW. Finally, we hope that the insights from this work will contribute to the development of more effective rehabilitation techniques.

## CONFLICT OF INTEREST STATEMENT

The authors declare no conflict of interest.

## Abbreviations

EEG: electroencephalogram
ERP: event-related-potentials
GFP: global field power
MRI: magnetic resonance imaging
sd: standard deviation
TW: travelling wave

## ACKNOWLEDGMENTS

The authors sincerely thank all participants of the naming experiment and the FNAF (Fédération Nationale des Aphasiques de France).

## Notes

### Competing Interest Statement

The authors have declared no competing interest.

## References

Aksenov, A; Beuter, A. (2021) Analysis of phase waves in the ECoG data. doi: 10.1051/mmnp/2021045

Ala-Salomäki, H; Kujala. J; Liljeström. M; Salmelin. R. (2021) Picture naming yields highly consistent cortical activation patterns: Test–retest reliability of magnetoencephalography recordings’. doi:10.1016/j.neuroimage.2020.117651.

Alamia, A.; VanRullen, R. (2019) Alpha oscillations and traveling waves: Signatures of predictive coding? doi:10.1371/journal.pbio.3000487.

Alario, F.X.; Ferrand, L. (1999) A set of 400 pictures standardized for French: Norms for name agreement, image agreement, familiarity, visual complexity, image variability, and age of acquisition, Behavior research methods, instruments, & computers, 31(3), pp. 531–552.

Alexander, D.M; Jurica. P; Trengove. C; Nikolaev. A.R; Gepshtein. S; Zvyagintsev. M, et al. (2013) Traveling waves and trial averaging: The nature of single-trial and averaged brain responses in large-scale cortical signals. doi:10.1016/j.neuroimage.2013.01.016.

Bessonov, N.; Beuter. A; Trofimchuk. S; Volpert. V. (2019) Cortical waves and post-stroke brain stimulation. doi:10.1002/mma.5620.

Bessonov, N; Beuter. A; Trofimchuk. S; Volpert. V (2020) Dynamics of periodicwaves in a neural field model. doi: 10.3390/math8071076.

Bhattacharya, S; Brincat. S.L; Lundqvist. M; Miller, E.K. (2022) Traveling waves in the prefrontal cortex during working memory. doi:10.1371/journal.pcbi.1009827.

Blumstein, S., Isaacs, E.; Mertus, J. (1982) The role of the gross spectral shape as a perceptual cue to place articulation in initial stop consonants. doi: 10.1121/1.2018824

Botella-Soler, V; Valderrama. M; Crépon. B; Navarro. V; Le Van Quyen. M. (2012) Large-scale cortical dynamics of sleep slow waves. doi:10.1371/journal.pone.0030757.

Brainard, D.H. (1997) The Psychophysics Toolbox. doi: 10.1163/156856897X00357

Caramazza, A. (1997). How Many Levels of Processing Are There in Lexical Access?. doi: 10.1080/026432997381664

Chantsoulis, M; Polrola. P; Goral-Polrola. J; Hajdukiewicz. A; Supinski. J; Kropotov. J.D, et al. (2017) Application of ERPs neuromarkers for assessment and treatment of a patient with chronic crossed aphasia after severe TBI and long-term coma – Case report. doi:10.5604/12321966.1232770.

Croce, P; Quercia. A; Costa. S; Zappasodi. F. (2020) EEG microstates associated with intra- and inter-subject alpha variability. doi: 10.1038/s41598-020-58787-w.

Davis, Z.W; Muller. L; Martinez-Trujillo. J; Sejnowski. T; Reynolds. J.H. (2020) Spontaneous travelling cortical waves gate perception in behaving primates. doi: 10.1038/s41586-020-2802-y.

Delorme, A; Makeig, S. (2004) EEGLAB: an open source toolbox for analysis of single-trial EEG dynamics including independent component analysis. doi: 10.1016/j.jneumeth.2003.10.009

Dell GS, Schwartz MF, Martin N, Saffran EM, Gagnon DA. (1997) Lexical access in aphasic and nonaphasic speakers. doi: 10.1037/0033-295x.104.4.801. PMID: 9337631.

Dell, G.S., Martin, N., Schwartz, M.F. (2007) A case-series test of the interactive two-step model of lexical access: Predicting word repetition from picture naming. doi: 10.1016/j.jml.2006.05.007

Derouesné, C; Poitreneau. J; Hugonot. L; Kalafat. M; Dubois. B; Laurent. B. (1999) Le Mental-State Examination (MMSE): un outil pratique pour l’évaluation de l’état cognitif des patients par le clinicien. Version française consensuelle. Presse Méd., 28, p.: 1141-8.

Devlin, J.T., Matthews, P.M., and Rushworth, M.F.S. (1998) Semantic Processing in the Left Inferior Prefrontal Cortex: A Combined Functional Magnetic Resonance Imaging and Transcranial Magnetic Stimulation Study, pp. 71–84. doi: 10.1162/089892903321107837

Donoghue, T; Haller. M; Peterson. E.J; Varma. P; Sebastian. P; Gao. R, et al. (2020) Parameterizing neural power spectra into periodic and aperiodic components. doi: 10.1038/s41593-020-00744-x.

Ermentrout, G. Bard; Kleinfeld, D. (2001) Traveling Electrical Waves in Cortex. doi: 10.1016/s0896-6273(01)00178-7.

Ermentrout, G Bard; Kleinfeld, D. (2001) Traveling Electrical Waves in Cortex: Insights from Phase Dynamics and Speculation on a Computational Role the results. doi: 10.1016/S0896-6273(01)00178-7

Fisher, N. (1993) Statistical Analysis of Circular Data. Cambridge: Cambridge University Press. [Preprint]. doi: 10.1017/CBO9780511564345.

Flamand-Roze, C; Falissard. B; Roze. E; Maintigneux. L; Beziz. J; Chacon. A, et al. (2011) Validation of a new language screening tool for patients with acute stroke: the Language Screening Test (LAST) doi:10.1161/STROKEAHA.110.609503.

Graves WW, Grabowski TJ, Mehta S, Gordon JK. (2007) A neural signature of phonological access: distinguishing the effects of word frequency from familiarity and length in overt picture naming. J Cogn Neurosci. doi: 10.1162/jocn.2007.19.4.617. PMID: 17381253

Griffin, Z and Bock, K. (1998). Processing Levels in Spoken Word Production. doi: 10.1006/jmla.1997.2547

Hampshire, A; Chamberlain. S.R; Monti. M.M; Duncan. J; Owen. A.M. (2010) The role of the right inferior frontal gyrus: inhibition and attentional control. doi:10.1016/j.neuroimage.2009.12.109.

Hartwigsen, G. (2015) The neurophysiology of language: Insights from non-invasive brain stimulation in the healthy human brain. doi:10.1016/j.bandl.2014.10.007.

Hartwigsen, G. (2016) Adaptive Plasticity in the Healthy Language Network: Implications for Language Recovery after Stroke. doi:10.1155/2016/9674790.

Hartwigsen, G; Stockert. A; Charpentier. L; Wawrzyniak. M; Klingbeil. J; Wrede. K, et al. (2020) Short-term modulation of the lesioned language network. doi:10.7554/eLife.54277.

Hartwigsen, G; Volz, L.J. (2021) Probing rapid network reorganization of motor and language functions via neuromodulation and neuroimaging. doi:10.1016/j.neuroimage.2020.117449.

Hassan, M; Benquet. P; Biraben. A; Berrou. C; Dufor. O; Wendling. F. (2015) Dynamic reorganization of functional brain networks during picture naming. doi:10.1016/j.cortex.2015.08.019.

Hemanth (2022) hausdorffDist (a,b), (https://www.mathworks.com/matlabcentral/fileexchange/56336-hausdorffdist-a-b), MATLAB Central File Exchange. Retrieved May 13, 2022.

Hodgson, V.J., Ralph, M.A.L; Jackson, R.L. (2021) NeuroImage Multiple dimensions underlying the functional organization of the language network. doi:10.1016/j.neuroimage.2021.118444.

Humphreys, G.W., Riddoch, M.J; Quinlan. P.T. (1998). Cascade processes in picture identification. doi: 10.1080/02643298808252927

Indefrey, P. (2011) The spatial and temporal signatures of word production components: A critical update. doi:10.3389/fpsyg.2011.00255.

Indefrey, P; Levelt, W.J.M. (2004) The spatial and temporal signatures of word production components. doi:10.1016/j.cognition.2002.06.001.

Kabbara, A; El Falou. W; Khalil. M; Wendling. F; Hassan. M. 52017) The dynamic functional core network of the human brain at rest. doi: 10.1038/s41598-017-03420-6.

Kalafat, M., Hugonot-Diener. L; Poitrenaud, J. (2003) Standardisation et étalonnage français du « Mini Mental State » (MMS) version GRECO., Rev Neuropsycol, 13(2), pp. 209–236.

Kamada, K; Saguer. M; Möller. M; Wicklow. K; Katenhäuser. M; Kober. H, et al. (1997) Functional and Metabolic. Analysis of Cerebral Ischiemia Using Magnetoenephalography and Proton Magnetic Resonance Spectroscopy. doi: 10.1002/ana.410420405

Klaus, J; Hartwigsen, G. (2019) Dissociating semantic and phonological contributions of the left inferior frontal gyrus to language production. doi:10.1002/hbm.24597.

Klimesch, W. (2018) The frequency architecture of brain and brain body oscillations: an analysis. doi:10.1111/ejn.14192.

Klimesch, W., Sauseng, P; Hanslmayr, S. (2007) EEG alpha oscillations: The inhibition-timing hypothesis. doi:10.1016/j.brainresrev.2006.06.003.

Klingbeil, J; Wawrzyniak. M; Stockert. A; Saur. D. (2019) Resting-state functional connectivity: An emerging method for the study of language networks in post-stroke aphasia. doi:10.1016/j.bandc.2017.08.005.

Kohn, S.E. (1988) Phonological Production Deficits in Aphasia. doi: 978-1-4615-7581-8_4.

Laganaro, M. (2017) Inter-study and inter-Individual Consistency and Variability of EEG/ERP Microstate Sequences in Referential Word Production. doi: 10.1007/s10548-017-0580-0.

Laganaro, M; Zimmermann, C. (2010) Origin of phoneme substitution and phoneme movement errors in aphasia, Language and Cognitive Processes. doi: 10.1080/01690960902719259

Lakatos, P; Shah. A.S; Knuth. K.H; Ulbert. I; Karmos. G; Schroeder. C.E. (2005) An Oscillatory Hierarchy Controlling Neuronal Excitabilityand Stimulus Processing in the Auditory Cortex. doi:10.1152/jn.00263.2005.

Lehmann, D; Skrandies, W. (1980) Reference-free identification of components of checkerboard-evoked multichannel potential fields. doi: 10.1016/0013-4694(80)90419-8.

Lehmann, D; Skrandies, W. (1984) Spatial Analysis of Evoked Potentials in Man-A review. doi: 10.1016/0301-0082(84)90003-0.

Levelt, W; Roelofs, A; Meyer. A.S. (1999). Multiple perspectives on word production. doi: 10.1017/S0140525X99451775

Liljeström, M; Kujala. J; Stevenson. C; Salmelin. R. (2015) Dynamic reconfiguration of the language network preceding onset of speech in picture naming. doi:10.1002/hbm.22697.

Lindvall, O; Kokaia, Z. (2015) Neurogenesis following stroke affecting the adult brain. doi:10.1101/cshperspect.a019034.

Llorens, A; Trébuchon. A; Liégois-Chauvel. C; Alario. F-X. (2011) Intra-cranial recordings of brain activity during language production. doi:10.3389/fpsyg.2011.00375ï.

Massimini, M; Huber. R; Ferrarelli. F; Hill. S; Tononi. G. (2004) The sleep slow oscillation as a traveling wave. doi:10.1523/JNEUROSCI.1318-04.2004.

Mehrkanoon, S; Breakspear. M; Britz. J; Boonstra. T.W. (2014) Intrinsic coupling modes in source-reconstructed electroencephalography. doi:10.1089/brain.2014.0280.

Meinzer, M; Elbert. T; Wienbruch. C; Djundja. D; Barthel. G; Rockstroch. B. (2004) Intensive language training enhances brain plasticity in chronic aphasia. doi: 10.1186/1741-7007-2-20.

Mheich, A; Hassan. M; Khalil. M; Berrou. C; Wendling. F. (2015) A new algorithm for spatiotemporal analysis of brain functional connectivity. doi:10.1016/j.jneumeth.2015.01.002.

Mheich, A; Dufor. O; Yassine. S; Kabbara. A; Biraben. A; Wendling. F, et al. (2021) HD-EEG for tracking sub-second brain dynamics during cognitive tasks. doi: 10.1038/s41597-021-00821-1.

Moliadze, V; Sierau. L; Lyzhko. E; Stenner. T; Werchowski. M; Siniatchkin. M, et al. (2019) After-effects of 10 Hz tACS over the prefrontal cortex on phonological word decisions. doi:10.1016/j.brs.2019.06.021.

Muller, L; Chavane. F; Reynolds. J; Sejnowski. T (2018a) Cortical travelling waves: mechanisms and computational principles. doi:10.1038/nrn.2018.20.

Murphy, M; Riedner. B.A; Huber. R; Massimini. M; Ferrarelli. F; Tononi. G. (2009) Source modeling sleep slow waves. doi:10.1073/pnas.0807933106.

Murray, M.M., Brunet, D; Michel, C.M. (2008) Topographic ERP analyses: A step-by-step tutorial review. doi: 10.1007/s10548-008-0054-5.

Oldfield, R.C. (1971) The assessment and analysis of handedness: The Edinburgh inventory. doi: 10.1016/0028-3932(71)90067-4.

Patten, T.M; Rennier. C.J; Robinson. P.A; Gong. P. (2012) Human cortical traveling waves: Dynamical properties and correlations with responses. doi:10.1371/journal.pone.0038392.

Petrovic, J; Milosevic. V; Zivkovic. M; Stojanov. D; Milojkovic. O; Kalauzi. A, et al. (2017) Slower EEG alpha generation, synchronization and “flow”-possible biomarkers of cognitive impairment and neuropathology of minor stroke. doi:10.7717/peerj.3839.

Pham, P; Roux. S; Matonti. F; Dupont. F; Agache. V; Chavane. F. (2013) Post-implantation impedance spectroscopy of subretinal micro-electrode arrays, OCT imaging and numerical simulation: Towards a more precise neuroprosthesis monitoring tool. doi:10.1088/1741-2560/10/4/046002.

Piai, V; Zheng, X. (2019) Psychology of Learning and Motivation, in Federmeier, K.D. (ed.) doi:10.1016/bs.plm.2019.07.002.

Price, C.J. (2000) The anatomy of language: contributions from functional neuroimaging, 44, pp. 335–359. doi: 10.1046/j.1469-7580.2000.19730335.x

Price, C.J. (2010) The anatomy of language: a review of 100 fMRI studies published in 2009. doi:10.1111/j.1749-6632.2010.05444.x.

Rapp, B; Goldrick, M. (2006). Speaking words: Contributions of cognitive neuropsychological research. doi: 10.1080/02643290542000049

Roelofs, A; Shitova, N. (2017) Importance of response time in assessing the cerebral dynamics of spoken word production: comment on Munding et al. (2016). doi:10.1080/23273798.2016.1274415.

Saha, S; Baumert, M. (2020) Intra- and Inter-subject Variability in EEG-Based Sensorimotor Brain Computer Interface: A Review. doi: 10.3389/fncom.2019.00087.

Salmelin, R; Hari. R; Lounasmaa. O.V; Sams. M. (1994) Dynamics of brain activation during picture naming. doi: 10.1038/368463a0.

Saravani, A. G; Forseth KJ; Tandon N; Pitkow X. (2019) Dynamic Brain Interactions during Picture Naming. doi: 10.1523/ENEURO.0472-18.2019.

Sarubbo, S; Tate. M; De Benedictis. A; Merler. S; Morits-Gasser. S; Herber. G, et al. (2020) Mapping critical cortical hubs and white matter pathways by direct electrical stimulation: an original functional atlas of the human brain. doi:10.1016/j.neuroimage.2019.116237.

Saur, D; Lange. R; Baumgaertner. A; Schraknepper. V; Willmes. K; Rijntjes. M, et al. (2006) Dynamics of language reorganization after stroke. doi:10.1093/brain/awl090.

Spironelli, C; Angrilli, A. (2009) EEG delta band as a marker of brain damage in aphasic patients after recovery of language. doi:10.1016/j.neuropsychologia.2008.10.019.

Sporns, O. (2013) The human connectome: Origins and challenges. doi:10.1016/j.neuroimage.2013.03.023.

Stockert, A; Wawrzyniak. M; Klingbeil. J; Wrede. K; Kümmerer. D; Hartwigsen. G, et al. (2020) Dynamics of language reorganization after left temporo-parietal and frontal stroke. doi:10.1093/brain/awaa023.

Strijkers, K., Costa, A; Thierry, G. (2010) Tracking lexical access in speech production: Electrophysiological correlates of word frequency and cognate effects. doi:10.1093/cercor/bhp153.

Van den Heuvel, M.P; Sporns, O. (2011) Rich-club organization of the human connectome, doi:10.1523/JNEUROSCI.3539-11.2011.

Vigneau, M; Beaucousin. V; Hervé. P-Y; Duffau. H; Crivello. F; Houdé. O, et al. (2006) Meta-analyzing left hemisphere language areas: Phonology, semantics, and sentence processing. doi:10.1016/j.neuroimage.2005.11.002.

Vigneau, M; Beaucousin. V; Hervé. P-Y; Jobard. G; Petit. L; Crivello. F, et al. (2011) NeuroImage What is right-hemisphere contribution to phonological, lexico-semantic, and sentence processing? Insights from a meta-analysis. doi:10.1016/j.neuroimage.2010.07.036.

Volpert, V.; Xu, B.; Tchechmedjiev, A.; Harispe, S.; Aksenov, A.; Mesnildrey, Q.; Beuter, A. (2022). Characterization of spatiotemporal dynamics in EEG data during picture naming with optical flow patterns. bioRxiv 2022.11.24.517789; doi: https://doi.org/10.1101/2022.11.24.517789

Zhang, H; Watrous. A; Patel. A; Jacobs. J. (2018) Theta and Alpha Oscillations Are Traveling Waves in the Human Neocortex. doi:10.1016/j.neuron.2018.05.019.

