## Supplementary Material S1 for "Investigating Spatiotemporal Dynamics of Cortical Activity During Language Production in the Healthy and Lesioned Brain"

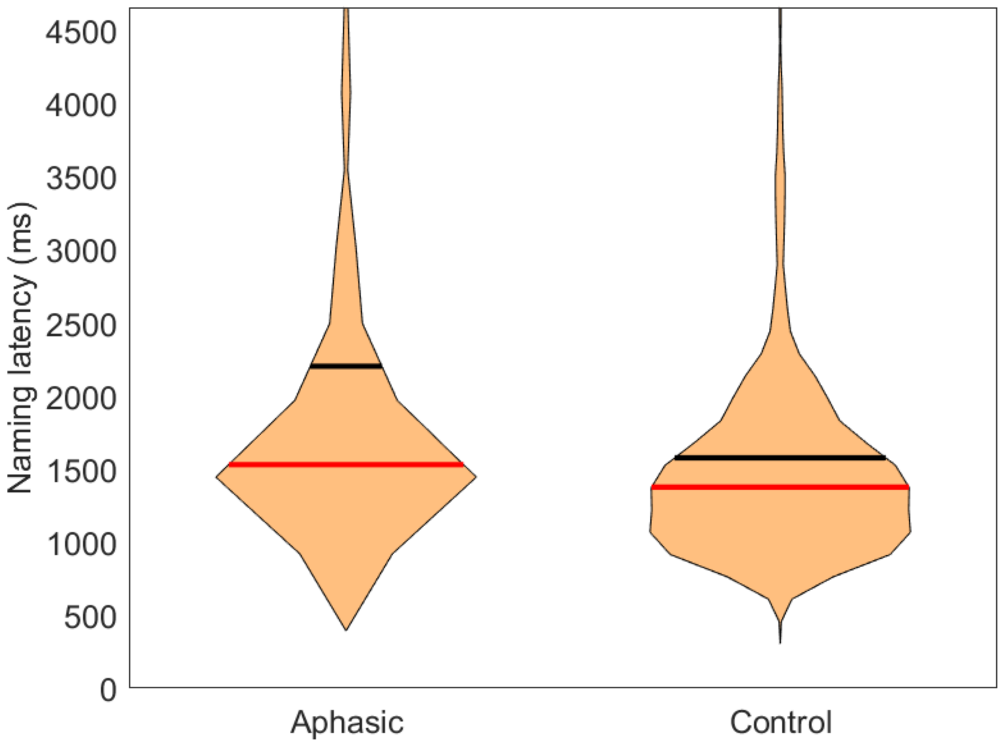


**S1:** Naming latency (in ms) of the aphasic and control groups. Red lines indicate the median latency, black lines indicate the mean latency.
