## Supplementary Material S2 for "Investigating Spatiotemporal Dynamics of Cortical Activity During Language Production in the Healthy and Lesioned Brain"

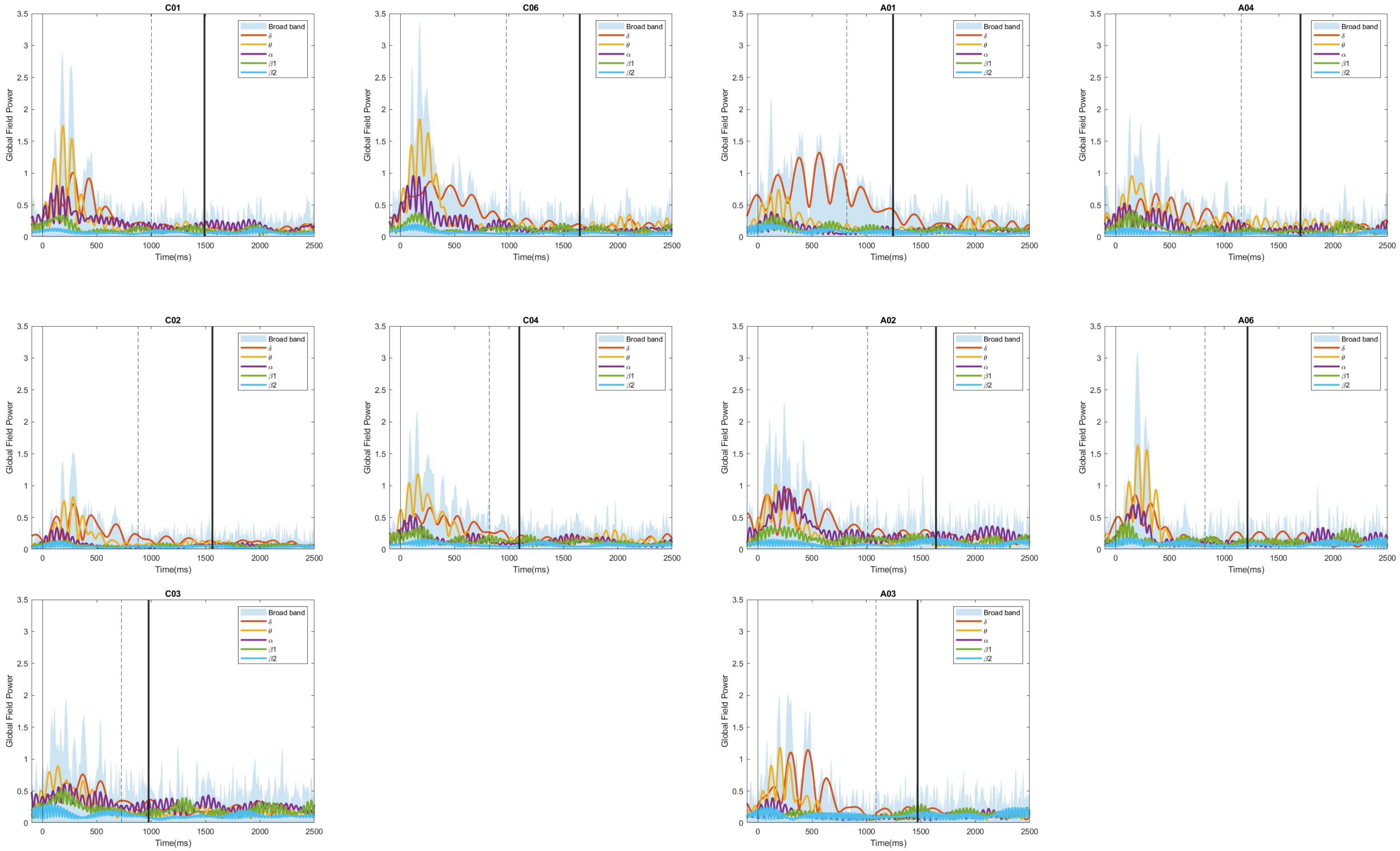


S2: Global field power for all ten subjects. In each panel, Blue area illustrates the broadband GFP and color lines illustrate the GFP in subranges.
