## Supplementary Material S3 for "Investigating Spatiotemporal Dynamics of Cortical Activity During Language Production in the Healthy and Lesioned Brain"

A. "In phase" ERP signals from C01 in the theta band

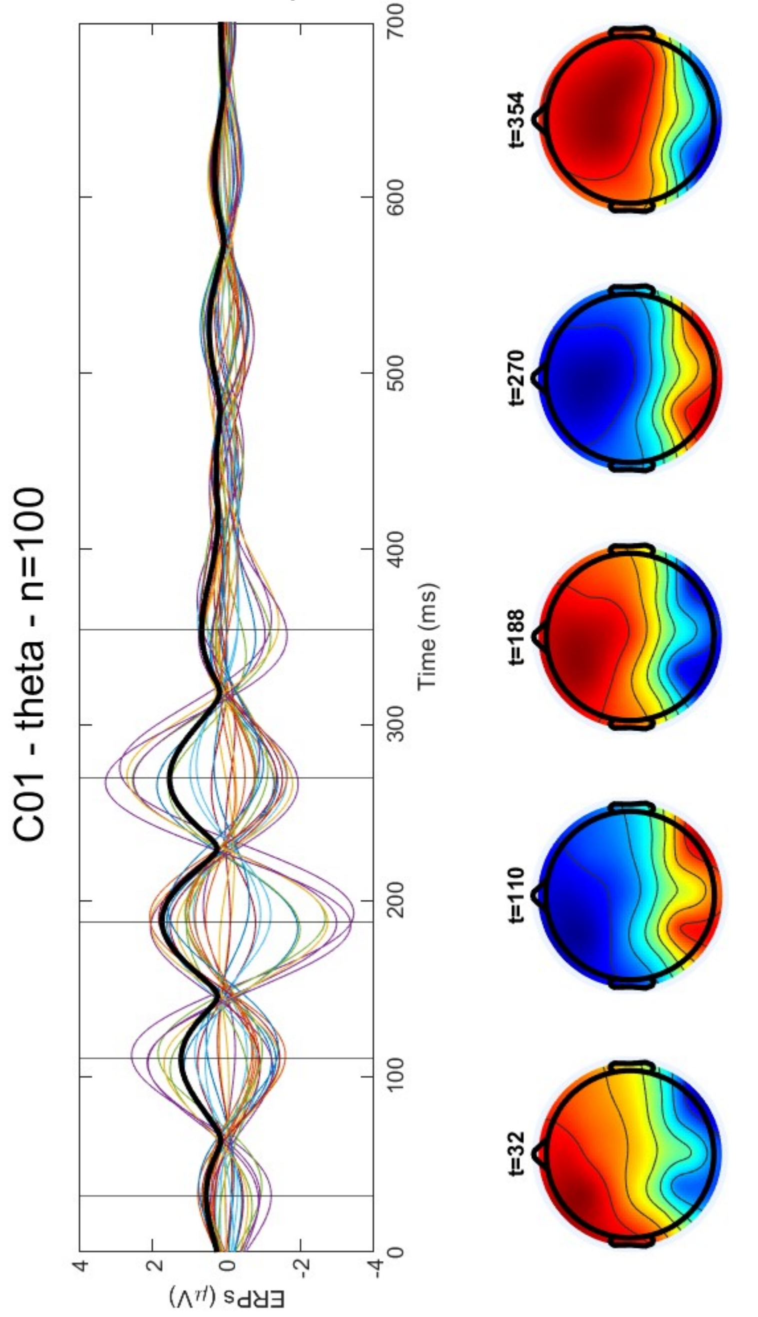


B. "Out of  phase" ERP signals from C04 in the theta band

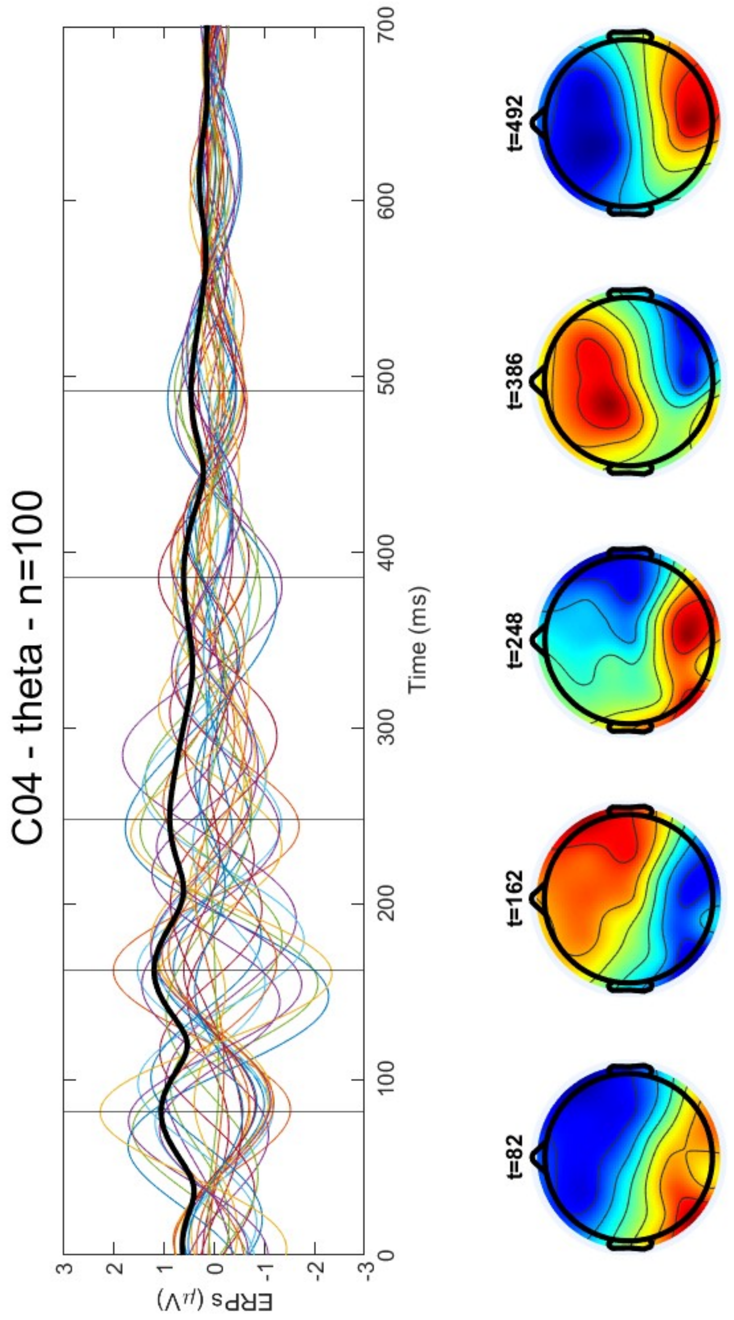


S3:
Example of in-phase and out of phase ERPs for subjects C01 (A) and C04 (B), in the theta range for all trials (n=100).
For each panel, colored lines illustrate the ERP signals, the black thick lines represent the GFP and topographic maps of activity are computed for the peaks of GFP illustrated by vertical lines.
