## Supplementary figures and images for "Investigating Spatiotemporal Dynamics of Cortical Activity During Language Production in the Healthy and Lesioned Brain"

### Supplementary Material S4

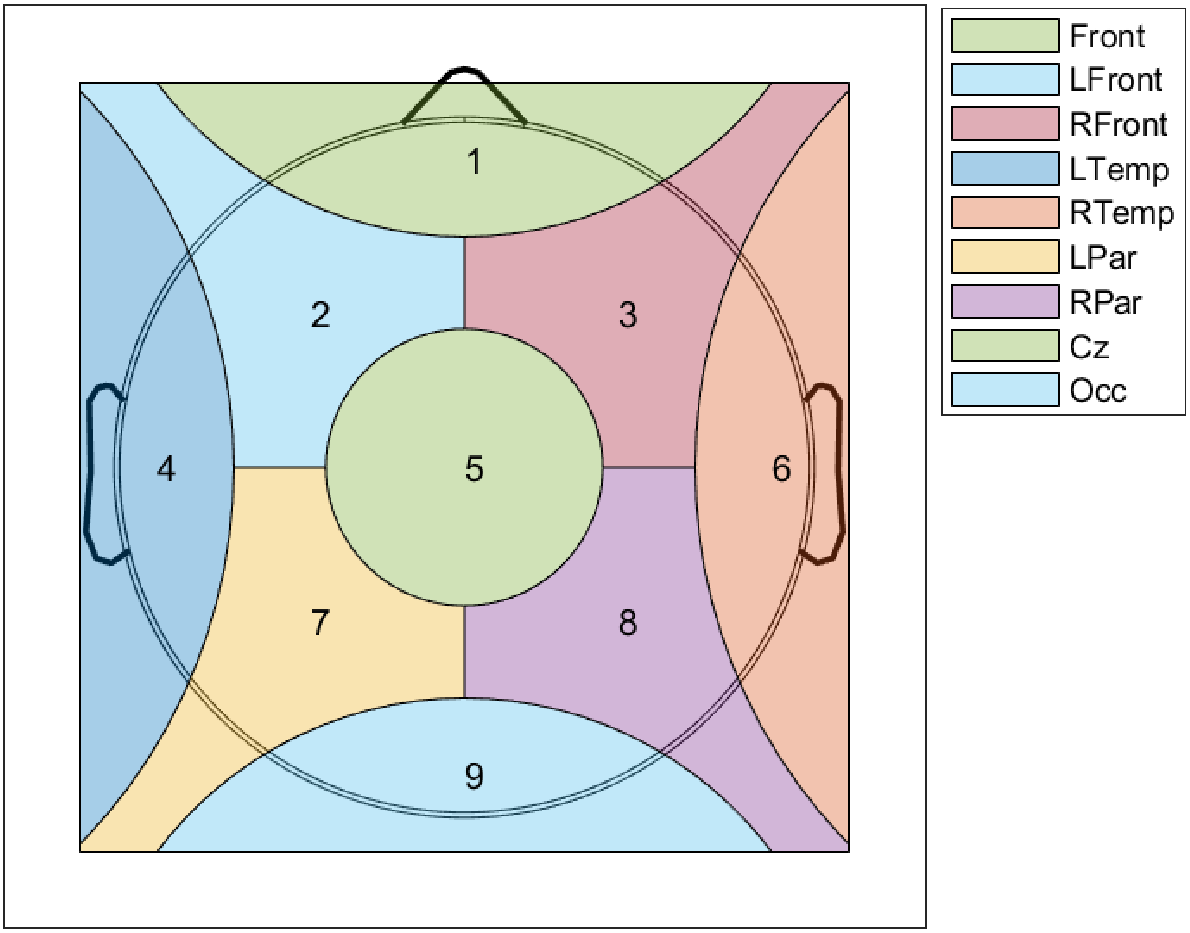


**S4:** Schematic view of the parcellation of the scalp surface used in figure 5.
