## Supplementary Material S5 for "Investigating Spatiotemporal Dynamics of Cortical Activity During Language Production in the Healthy and Lesioned Brain"

**S5:** Example of the distribution of amplitude waves detected at 22Hz for each subject in ten repetition of the control word “Chien” (i.e., “Dog”).

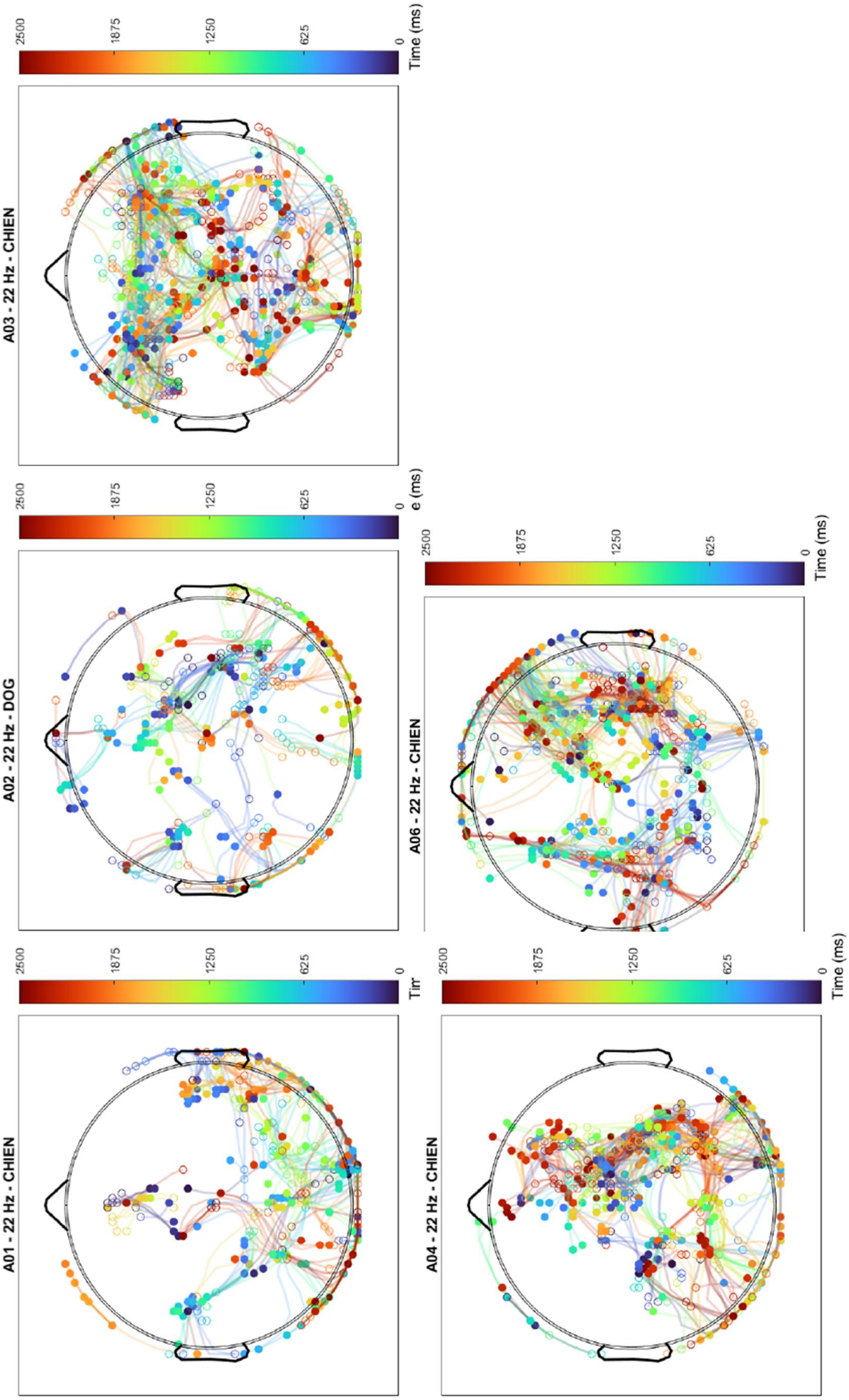

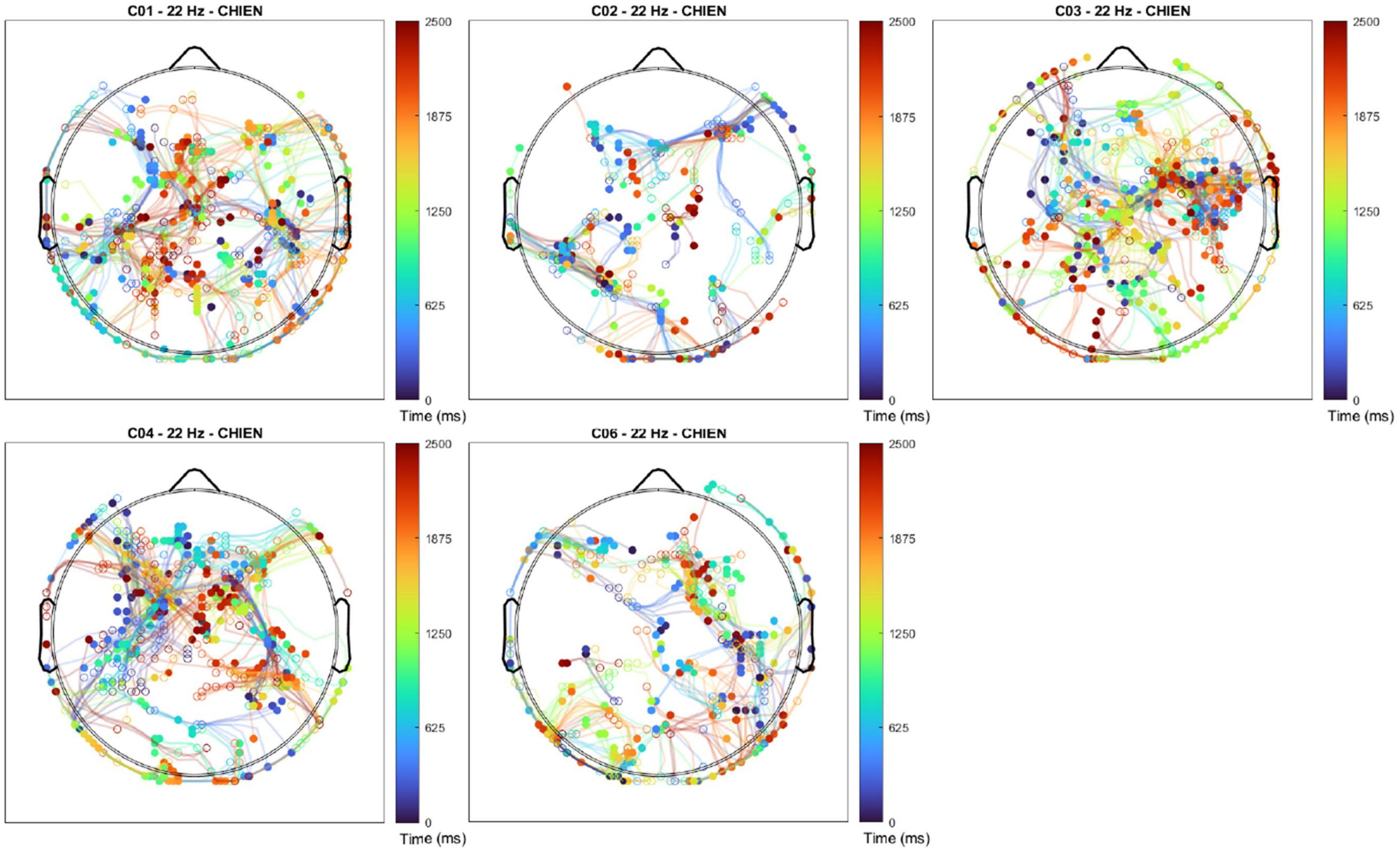
C01 - 22 Hz - CHIEN
C02 - 22 Hz - CHIEN
C03 - 22 Hz - CHIEN
C04 - 22 Hz - CHIEN
C06 - 22 Hz - CHIEN
2500
1875
1250
625
0
Time (ms)
